# Singular Odorant Receptor Expression Orchestrated by Promoter Activation Specificity in *Apis Mellifera* Olfactory Sensory Neurons

**DOI:** 10.1101/2024.05.04.592513

**Authors:** Weixing Zhang, Yage Nie, Yiheng Li, Yicong Xu, Tao Xu, Peiyu Shi, Fang Liu, Hongxia Zhao, Qing Ma, Jin Xu

## Abstract

Honeybee, positioned as an outgroup of flies and mosquitos, have exhibited intriguing duplications and expansions of OR genes in genomic studies. However, little is known about how the OR genes are expressed and regulated in honeybee olfactory sensory neurons (OSNs). In this study, we utilized a single-cell multi-omics approach to profile the transcriptome and chromatin accessibility in *Apis mellifera* antennal nuclei, aiming to elucidate OR gene expression and its underlying regulatory mechanisms. Our systematical analysis unveiled a similar singular expression pattern of ligand-specific receptors in *Apis mellifera* OSNs, parallel with those observed in *Drosophila melanogaster*. Mechanistically, we discovered that promoter activation of OR genes orchestrates receptor expression patterns. For instance, although multiple adjacent OR genes are co-expressed with a single active promoter through polycistronic transcription, only the first OR gene could produce functional receptor protein, as supported by transcriptome quantification in OSNs. Additionally, we found that co-expression of receptor proteins might occur only when co-expressed OR genes possess multiple accessible promoters. This scenario is less common than expected, considering the number and evolutionary age of OR genes in *Apis mellifera*, suggesting a selection favoring the specialization of rapidly expanded OR genes in *Apis mellifera*. Overall, our study provides significant insights into the insect olfaction system and the regulatory mechanisms of OR genes expression.

## Introduction

The olfactory sense in insects is critical for their ability to detect and discriminate a wide range of volatile chemicals in their surroundings. This chemosensory detection is facilitated by receptors encoded by three major families: gustatory receptors (GRs), ionotropic receptors (IRs), and odorant receptors (ORs) ^1,2^. Among these, ORs play a predominant role in detecting chemically diverse odors in most species. ORs function as odorant-gated ion channels ^3,4^ formed by a ligand-specific receptor and a heteromultimeric complex of the conserved odorant receptor co-receptor (*Orco*) ^5,6^. Studies in the model organism *Drosophila melanogaster* (*D. melanogaster*) have elucidated key principles of OR gene expression, in where each OSN expresses a single ligand-specific OR gene ^7–9^. OSNs expressing the same ligand-specific OR gene share the identical cellular features and chemosensory function, defining an OSN class. The "one receptor to one neuron" principle, which succinctly explains olfaction specificity, has been widely accepted and is supported by extensive studies on vertebrate olfactory systems ^10–15^. In fact, in *Drosophila*, three out of the 34 OSN classes in the antenna also express two or three different ligand-specific OR genes according to systematic assays ^16^. Studies in *Anopheles gambiae*, have also shown co-expression of six adjacent OR genes (*AgOr47*, *AgOr16*, *AgOr17*, *AgOr13*, *AgOr55*, and *AgOr15*) as detected by fluorescence *in situ* hybridization ^17,18^. These previous reported co-expressed OR genes are tandem duplications, located adjacently in the genome and exhibiting higher sequence similarity. The occasional co-expression of OR genes is explained as representing an early evolutionary stage of duplicated gene pairs, as newly duplicated genes tend to maintain the same expression pattern as their ancestry gene ^19,20^.

However, recent investigations in non-model insect species have challenged this foundational principle ^16,17,21,22^. Single-cell transcriptome study in *Aedes aegypti* (*Ae. aegypti*) has highlighted genome-wide co-expression of multiple divergent chemoreceptors in OSNs ^23^. The absence of a molecular mechanism to regulate the exclusive expression of individual OR gene, as elucidated in the vertebrate olfactory system ^24–27^, suggests that co-expression of multiple OR genes within a single OSN is plausible occurrence in insects. This raises the question of which form is the norm and preferred across different insect orders.

Genome analysis reveals a remarkable expansion of the OR genes family over varying evolutionary timescale in *Apis mellifera* (*A. mellifera*), providing an intriguing system to investigate. The *A. mellifera* genome contains more than 160 OR genes, some of which exhibit high sequence divergence, while others display nearly identical sequences due to recent duplication events ^28^. Recent advancements in single-cell multi-omics approaches have provided powerful tools for investigating OR genes expression specificity and OSN cellular divergence in non-model species. In this study, we conducted a comprehensive cell census with simultaneously profiling gene expression and chromatin accessibility of *A. mellifera* worker antenna tissues for the first time. Our aim is to characterize the expression patterns of OR genes in *A. mellifera* OSNs, depict how OR gene expression is connected to OSN cellular identity, and investigate how the co-expression of multiple ligand-specific OR genes is tied to their evolutionary timescales.

## Results

### Single-cell profiling reveals abundance of olfactory sensory neurons in *A. mellifera* antennae

To examine the expression of chemosensory receptors at the single-cell level, we conducted single-cell multi-omics sequencing on antenna tissue from adult worker bees at three time points following emergence from the pupal stage (**Figure 1A, Figure S1A-C and Methods**). Employing unsupervised clustering, we classified the profiles from 8868 high-quality cells into six cell types and annotated them as neuron (*syt1*), epithelial (*grh*), glial (*repo*), sheath (*rpod*), and two supporting cell types (*Obp4* or *Obp5*), based on their expression and accessibility of conserved marker genes defined in *D. melanogaster* (**Figure 1B-C, Figure S1D**) ^29^. Additionally, we identified cell type-specific genes for each class, along with their corresponding functions, further supporting their cell type identities (**Figure S1E-F**).

**Figure 1.**
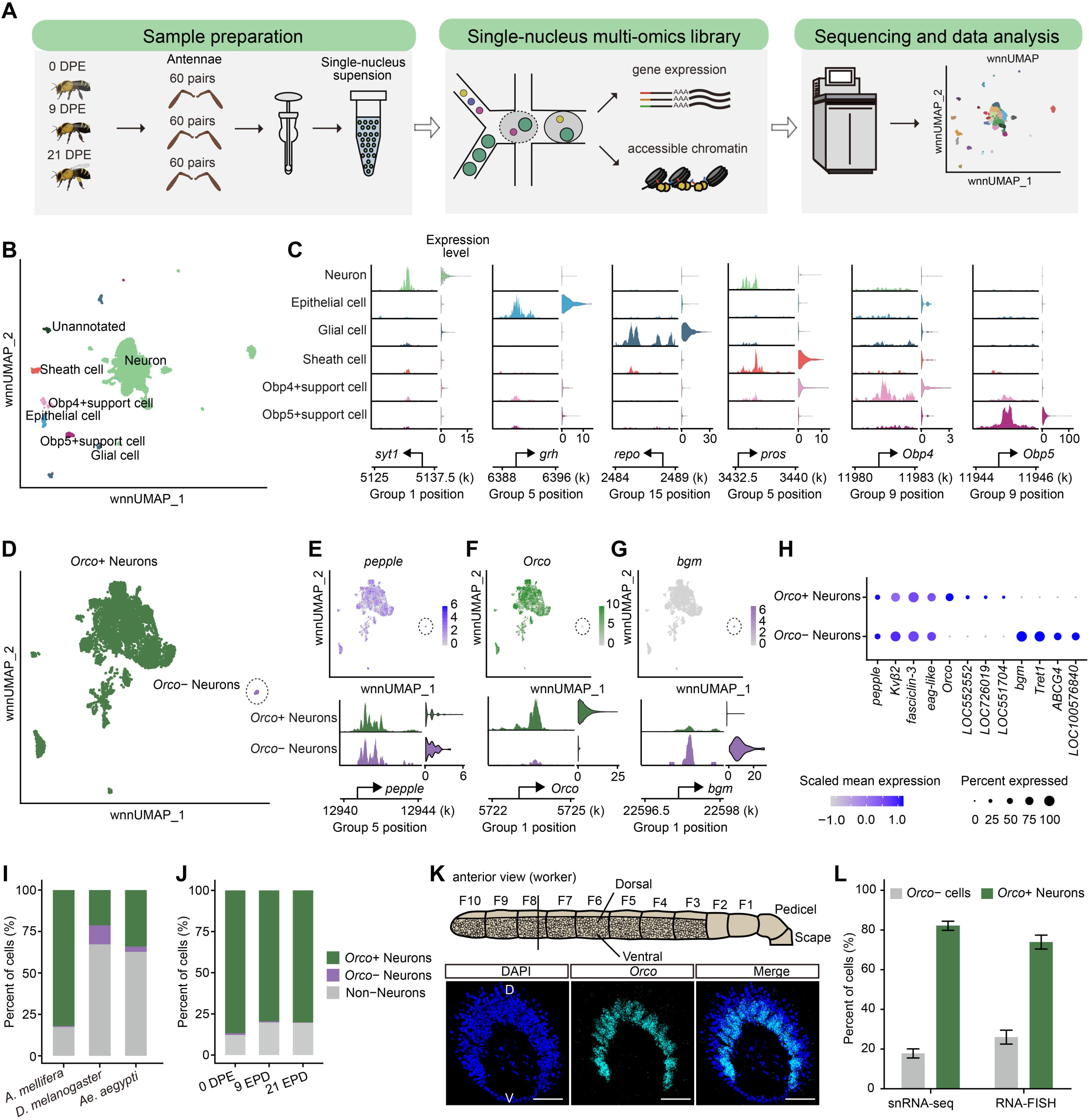
Cell census of *A. mellifera* worker antenna by single-cell multi-omics profiling (A) Schematic of the experimental design for single-nucleus multi-omics sequencing workflow. DPE: days post-eclosion. (B) UMAP of weighted-nearest neighbor (wnnUMAP) graph illustrating all *A. mellifera* antennal cells passing quality control, colored by annotated clusters. Cells that could not be defined to any cell types are unannotated. (C) Genome tracks illustrating the normalized chromatin accessibility surrounding marker genes. Next to the genome track is a violin plot representing the expression levels of selected marker genes in each major cell type. The genomic location of the marker genes is shown below. (D) The wnnUMAP plot displays all *A. mellifera* antennal neurons, with identified *Orco*+ and *Orco*– neuron clusters labeled in different colors. The dashed circle represents the *Orco*– neuron cells. (E-G) Expression mapped onto wnnUMAP plots for *pepple* as a marker for neuron cell type (E), *Orco* as a marker for *Orco*+ neuron cell type (F), and *bgm* as a marker for *Orco*– neuron cell type (G). The dashed circles represent *Orco*– neuron cells. Below the wnnUMAP, the genome track shows the normalized chromatin accessibility signal around the marker genes, along with a violin plot displaying the expression level of the marker genes in the *Orco*+ neuron or *Orco*– neuron cell type. The genomic location of the marker genes is shown below. (H) The bubble plot illustrating the expression of selected marker genes for *Orco*+ and *Orco*– neuron cell types. The color scale represents the scaled mean expression level across the *Orco*+ or *Orco*– neurons, while the size of the dots indicates the percentage of cells in each cell type expressing the marker gene. (I) The stacked bar plots depict the percentages of *Orco*+ neurons, *Orco*– neurons, and non-neuronal cell types in the antennae of *A. mellifera*, *D. melanogaster*, and *Ae. aegypti*. (J) The stacked bar plots display the percentages of *Orco*+ neurons, *Orco*– neurons, and non-neuronal cell types in *A. mellifera* antennae of different days post-eclosion. (K) The distribution of *Orco*-expressing cells on transversal sections through the antenna of *A. mellifera* workers is shown. The top panel presents a schematic representation of the *A. mellifera* antennae, showing distribution patterns of sensilla (redrawn from Esslen and Kaissling, 1976) ^58^. Olfactory sensilla are distributed from the third flagellar segment (F3) to the tenth flagellar segment (F10). The bottom panel displays high magnification images of transversal sections obtained through RNA-FISH with *Orco* specific probes. The images represent projections of z-stacks of confocal images and represent the results of three independent experiments using antennae from different workers. Scale bars: 50 µm. (L) The bar plots illustrate the percentages of *Orco*+/*Orco*– cells, as determined by snRNA-seq or RNA-FISH (K). The data represent the mean ± SEM, with n = 3.

Remarkably, we found that neurons constituted the most abundant cell type in *A. mellifera* antennae (**Figure S1G and Figure 1D**). These neurons were subsequently classified into two populations based on their expression of *Or2* (**Figure 1E-H**), the ortholog of *Orco* in *D. Melanogaster* ^30^. We observed that approximately 83% of cells in *A. mellifera* antennae were *Orco*+ neurons, which we designated as OSNs. This proportion is much higher than that observed in *D. Melanogaster* (∼33%) ^31^ and *Ae. Aegypti* (∼37%) ^32^ (**Figure 1I**). *A. mellifera* sensilla typically contain 18-35 OSNs ^33^, whereas *D. Melanogaster* ^34^ and *Ae. Aegypti* ^35^ sensilla contain 1-4 OSNs. The higher abundance of OSNs in *A. mellifera* antenna was expected, but accurate estimation was lacking. We further validated and quantified the proportion of *Orco*+ OSNs by RNA fluorescence *in situ* Hybridization (RNA-FISH) for *AmOrco* (**Figure 1K-L**), confirming the *A. mellifera* specific OSNs composition in antenna tissues. Additionally, there was no significant difference in cellular composition among *A. mellifera* at different days post-eclosion, suggesting the completion of antenna tissue development in adult worker of *A. mellifera* (**Figure 1J**). In brief, we have presented a high-quality single-cell multi-omics profile of *A. mellifera* antenna tissue and provided a comprehensive single-cell census.

### Discrepancy between OR gene expression and promoter accessibility

In *Drosophila*, the cellular identities of OSNs, as revealed by their transcriptome, is closely tied to the specific receptor they express. OSNs expressing the same OR gene tend to exhibit similar transcriptome profile and cluster together based on distinct transcription features ^36–38^. To further explore OSN classes in *A. mellifera*, we initially classified them into 46 sub-clusters using conventional unsupervised methodologies. However, we noticed that cells in the same sub-cluster still displayed heterogeneity, suggesting incomplete separation. To address this, we further employed an Iterative Subcluster After Clustering (ISAC) algorithm ^39^, which partitioned cells into subpopulations based on their transcriptomic distances, iteratively repeating this process until heterogeneity within subpopulations was eliminated (**Figure S2A-C, see Methods**). This approach yielded 115 distinct OSN classes, of which 59 with high discriminatory power were selected for further analysis (**Figure 2A-B, see Methods**).

**Figure 2.**
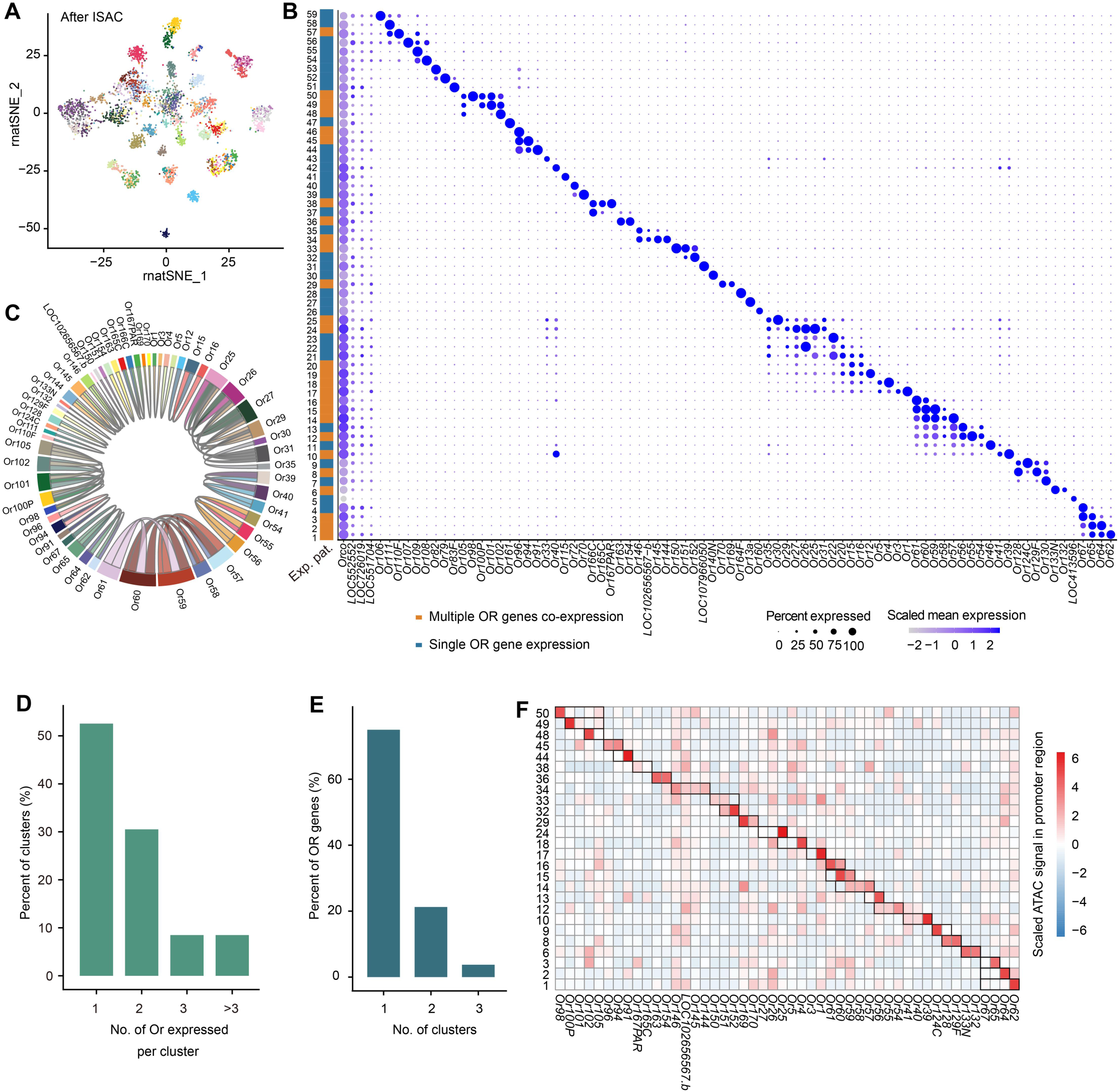
Cellular divergence and receptor expression patterns in *A. mellifera* OSNs. (A) The visualization of *Orco*+ neurons is accomplished using tSNE based on transcriptomic distance utilizing Iterative Subcluster After Clustering (ISAC). Cells are arranged based on transcriptome similarity. Each dot represents a cell. (B) The bubble plot displaying the expression of OR genes in each cell type. The size of the dot indicates the proportion of cells in that cell type expressing OR genes. The scaled mean expression of OR genes is represented from light blue to dark blue. (C) Chord plot shows the co-expressed pairs of OR genes that meet co-expression criteria for exhibiting significant OR genes expression: pct.exp > 40% and average expression scaled ≥ 2.5. (D) The bar plot shows the percentages of clusters expressing varying number of OR genes in *A. mellifera*. (E) The bar plot shows the percentages of OR genes expressed in either single or multiple clusters. (F) The heatmap illustrates the mean ATAC signal in the promoter regions of OR genes within co-expression clusters. Rows have been scaled. Black boxes indicate expressed OR genes within each cluster.

We then assessed the expression of OR genes in each OSN class and found that while OR genes demonstrated expression specificity within OSN classes, many OSN clusters exhibited co-expression of multiple OR genes (**Figure 2C**). Specifically, 47% of the OSN classes displayed expression of more than one ligand-specific OR genes (**Figure 2D**), and 25% OR genes were expressed in more than one OSN classes (**Figure 2E**). Interestingly, we found unexpected promoter accessibility patterns of co-expressed OR genes, as revealed by ATAC-seq profile from the same cells in each class. In the majority of OSN classes with co-expressed OR genes, only a single promoter was accessible (**Figure 2F**), prompting further investigation into these unexpected observations.

### Polycistronic transcription of adjacent OR genes via a single active promoter

To further elucidate these unexpected observations, we focused on OSN classes that exhibited co-expression of OR genes with a single accessible promoter (Co-expression (SP)) (**Figure 2F**). For instance, in cluster 24, co-expression of *Or25*, *Or26*, and *Or27* were observed and confirmed by RNA-FISH (**Figure 3A-C**). Interestingly, we found significant accessibility only in the promoter region of *Or25*, which is the upstream gene according to their transcription direction (**Figure 3D**). Similar chromatin accessibility and expression profiles for other typical OR genes were demonstrated in **Figure S3**. Since the co-expression was confirmed by an independent method, ruling out potential false discovery, we hypothesized that the downstream OR genes may undergo transcription as a single unit initiated from the accessible promoter. To test this hypothesis, we designed primer pairs to amplify the intergenic regions between OR genes using RT-PCR, which confirmed the occurrence of transcription readthrough events (**Figure 3E**). Taken together, these findings suggest that the co-expressed OR genes with a single active promoter are likely transcribed as polycistronic transcripts.

**Figure 3.**
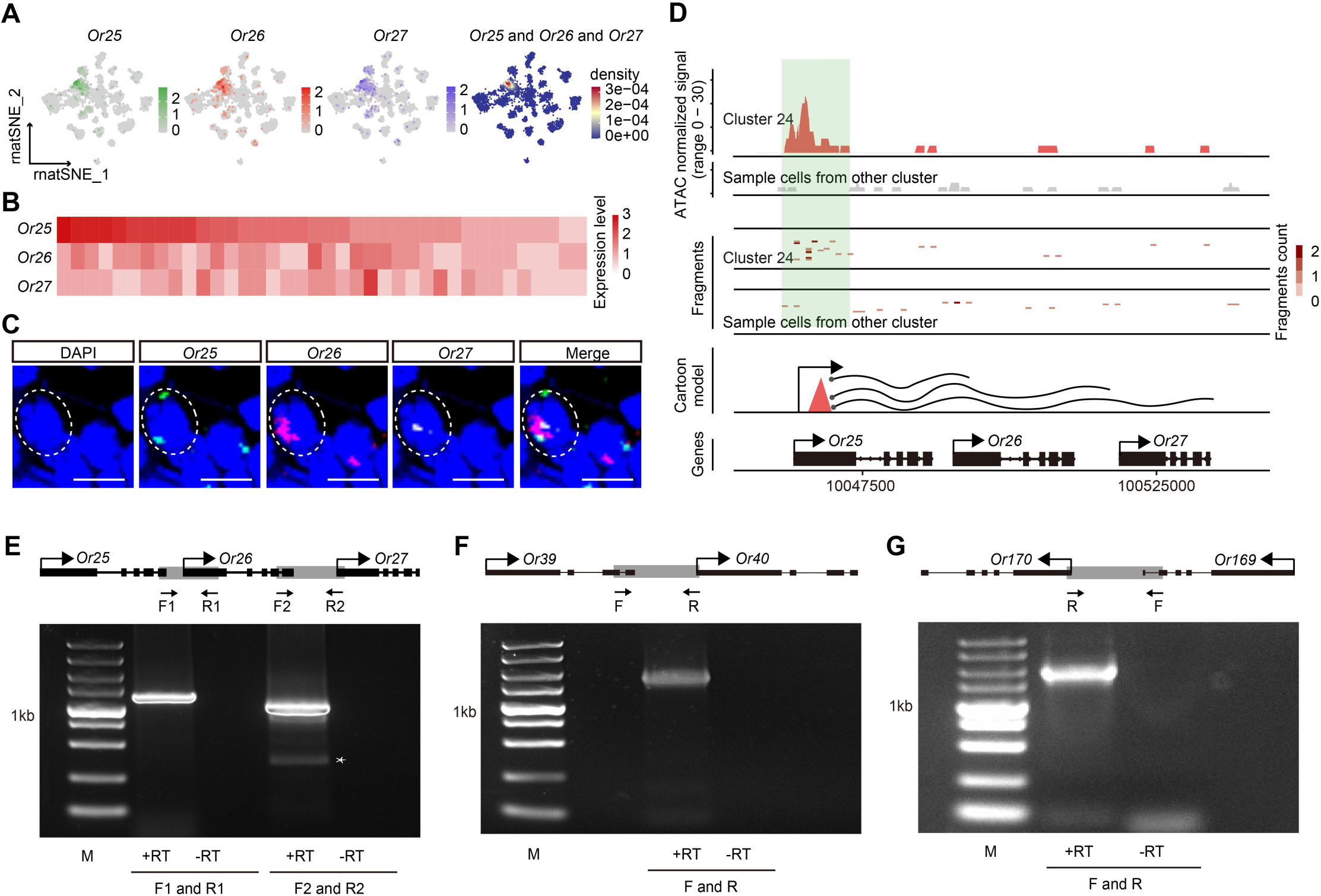
Co-expression of adjacent OR genes via polycistronic transcription in *A. mellifera* OSNs (A) The t-SNE plots visualize the expression of *Or25*, *Or26*, or Or27 in clusters. The co-expression t-SNE plots on the right depict the density of each single nucleus for *Or25*, *Or26*, and *Or27*. (B) The heatmaps display the co-expression of *Or25*, *Or26*, and *Or27* in a cluster. Rows represent receptors, and columns represent cells. The color of the heatmap represents the normalized expression (z-score). (C) High magnification images of a three-channel RNA-FISH experiment conducted on a transversal section of worker antenna. The section was incubated with antisense RNA probes for *Or25* (green), *Or26* (red) and *Or27* (cyan). The section was counterstained with DAPI (shown in blue). The dashed circles represent the nuclei of the neurons. Scale bars: 5 µm. (D) The genome tracks show the cell type-specific chromatin accessibility around *Or25*, *Or26*, and *Or27* in snATAC-seq data, with the promoter region highlighted in light green. In the fragments model, the tracks have been normalized based on the number of fragments in each cluster. The cartoon model represents OR gene expression patterns in cluster 24. Red triangles indicate the promoters of OR genes expressed in cluster 24. The wavy lines represent transcribed RNA molecules, and the solid circles represent the 5’ cap. In gene models, black boxes and thin solid lines represent exons and introns, respectively. The position and relative size of exons and introns were adopted from *A. mellifera* genome annotation and drawn to scale. The positions of the arrows represent the transcription start sites, while the direction of the arrows indicates the direction of transcription. (E-G) Agarose gel shows comparable amounts of products from RT-PCR analysis of nuclear RNA with a forward primer (F) located just upstream of the OR gene exon combined with a reverse primer (R) located downstream of the OR gene exon, as schematized in the top panel, *Or25*, *Or26*, and *Or27* (E), *Or39* and *Or40* (F), *Or169* and *Or170* (G). Asterisks indicate an unspecific band. ‘M’ indicates 1 kb molecular marker (Takara, 3428A).

### Cis-regulatory elements orchestrating co-expression OR genes with multiple active promoters

Besides the co-expression of OR genes via polycistronic transcription, we identified 5 OSN classes exhibiting co-accessibility at the co-expressed OR gene promoters (Co-expression (MP)). For instance, in cluster 8, the co-expression of *Or128* and *Or129F* within individual cells was validated by RNA-FISH (**Figure 4A-C**) and co-accessibility at the promoter region of *Or128* and *Or129F* was observed (**Figure 4D**). Similar patterns were also observed for other co-expression OR genes, such as the co-expression of *Or163* and *Or154*, as demonstrated in **Figure S4A-D**.

**Figure 4.**
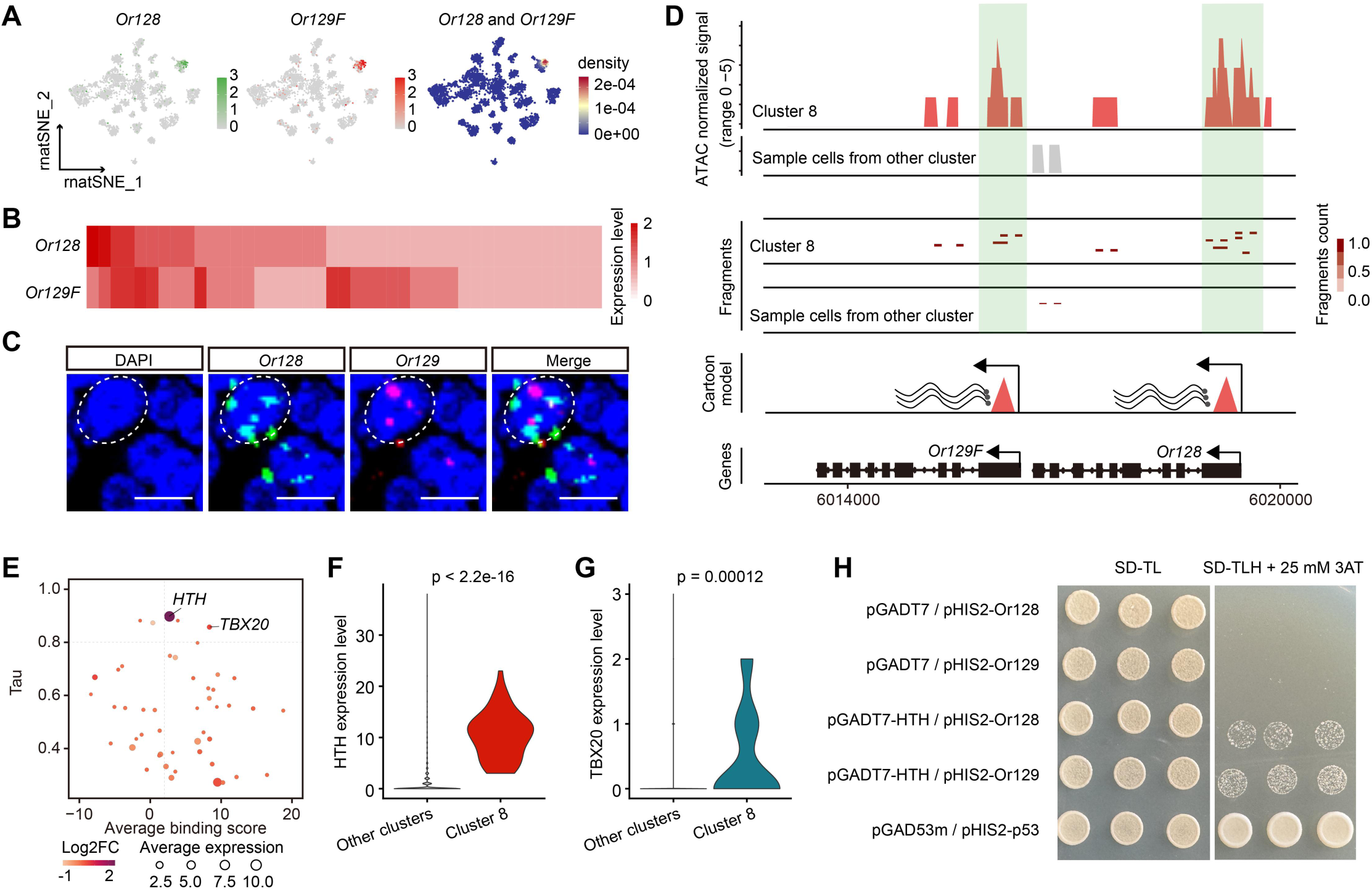
Co-expression of OR genes with multiple accessible promoters (A) The t-SNE plots visualize the expression of *Or128* (left) or *Or129F* (middle) in clusters. The density distribution of cells co-expressing *Or128* and *Or129F* is shown in the t-SNE plot on the right. (B) The heatmap demonstrates the co-expression of *Or128* and *Or129F* in the cluster. Rows indicate OR genes, and columns represent cells. The color of the heatmap represents the normalized expression. (C) High magnification images of a two-channel RNA-FISH experiment conducted on a transversal section of a worker antenna. The section was incubated with antisense RNA probes targeting *Or128* (green) and *Or129F* (red). DAPI staining (blue) was used as a counterstain. The dashed circles represent the nuclei of the neuron. Scale bars: 5 µm. (D) The genome tracks display the cell type-specific chromatin accessibility surrounding *Or128* and *Or129F* in snATAC-seq data. The promoter region is highlighted in light green. In the fragments model, the tracks have been normalized based on the number of fragments in each cluster. Multiple promoters refer to the promoter regions where the number of reads is greater than 10, and the fold change between the signals of multiple promoter regions is less than 2. The cartoon model depicts OR gene expression patterns in cluster 8. Red triangles indicate the promoters of OR genes expressed in cluster 8. The wavy lines represent transcribed RNA molecules, and the solid circles represent the 5’ cap. In the gene model, black boxes and thin solid lines represent exons and introns, respectively. The positions and relative sizes of exons and introns were obtained from *A. mellifera* genome annotation and drawn to scale. The positions of the arrows indicate the transcription start sites, while the direction of the arrows indicates the direction of transcription. (E) The bubble plot visualizes candidate TFs based on their characteristics. The Y-axis represents the Tau score, indicating expression specificity (higher values indicate higher specificity), while the X-axis depicts the Bindscore, an average measure of sequence matching between shared motif sequences and promoter regions (higher values indicate a stronger match). The size of the bubble represents the average expression of TFs, the color represents Log2FC of TF, and comparisons are made between clusters expressing *Or128* and *Or129F* and other clusters. (F-G) Violin plots display the expression of TFs *HTH* (F) and *TBX20* (G) from snRNA-seq data. The expression levels of TFs between cluster 8 and other clusters are compared by the Wilcoxon Rank Sum test, and the Bonferroni *p*-value is annotated above. (H) A Y1H assay was used to test the binding of the HTH transcription factor to the *Or128* and *Or129F* promoters. pGADT7 / pHIS2-Or128, pGADT7 / pHIS2-Or129, and pGAD53m / pHIS2-p53 vector were used as control.

In *D. melanogaster*, temporospatial-specific expression of transcription regulators (TFs) has been confirmed to be responsible for the specific expression of OR genes ^16,40,41^. If co-expression OR genes is programmed simultaneously, we would expect to observe higher promoter sequence similarity and the identification of common TF binding sites in the promoter regions of co-expressed OR genes, as previously demonstrated in DmelOr33c and DmelOr85e ^42,43^. Utilizing the TOBIAS ^44^ to identify TF binding sites within the open accessible regions (**Methods**), we found shared binding sites corresponding to homeobox protein (HTH) and the T-box transcription factor TBX20 (TBX20) at both *Or128* and *Or129F* promoter regions (**Figure 4E**). Conventionally, HTH and TBX20, which are conserved transcriptional activators in the nervous system ^37,45^, exhibited significant expression specificity in cluster 8, as depicted in **Figure 4F-G**. Yeast one-hybrid (Y1H) assays confirmed that HTH could recognize and bind to the promoter of *Or128* or *Or129F* (**Figure 4H**). Collectively, we demonstrated that the co-expression of OR genes with significant accessibility in the promoter in *A. mellifera* OSNs might be driven by common regulatory elements, following the same regulatory mechanism as proposed in *D. melanogaster*.

### Promoter activation of OR genes embedding the OSN cellular identity

More intriguing instances of polycistronic transcription and co-expression of OR genes were observed in a tandem duplication block, comprising *Or62*, *Or64*, *Or65*, and *Or67* (**Figure 5A**). Integrating the expression and chromatin accessibility profiles, we revealed that the promoters of these four OR genes were exclusively open, and the accessible promoter tended to facilitate polycistronic transcription of the downstream genes (**Figure 5B**). For instance, in the OSNs with an accessible promoter at *Or62*, expression of *Or62*, *Or64*, *Or65*, *Or67* was observed. Notably, there was a dominance of expression of 5′-located OR genes over 3′-located genes (**Figure 5A-B**), resulting in a differential expression of down-stream genes. Similarly, in OSNs with an accessible promoter at *Or64*, transcription of the downstream OR genes was observed, but no expression of *Or62* (the upstream OR genes) (**Figure 5A-B**). This pattern was consistent across cluster 3 and cluster 4 (**Figure 5A-B**). The expression of the four OR genes was further validated by RNA-FISH (**Figure S5A-C**), excluding the possibility of contaminations during single cell library construction.

**Figure 5.**
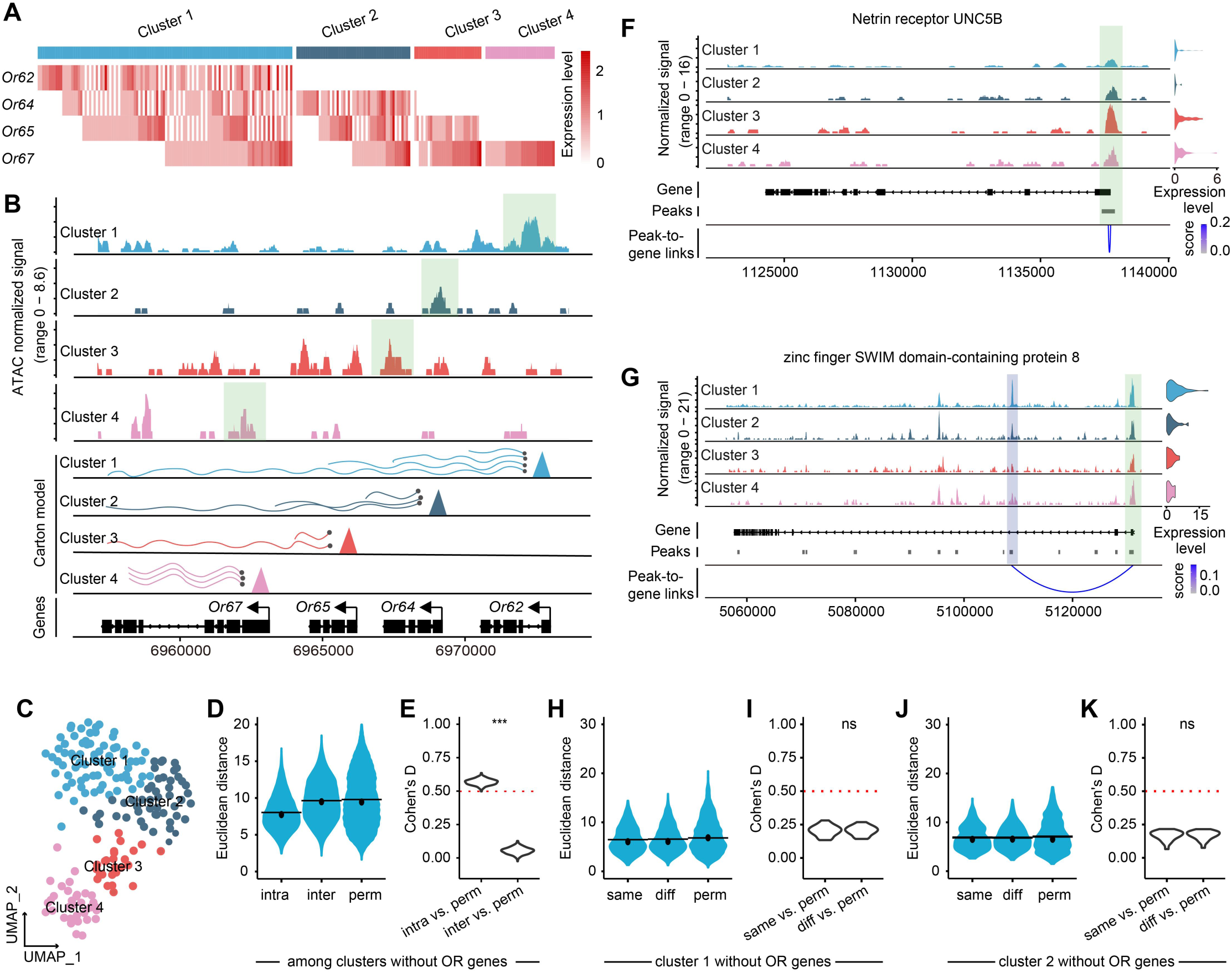
Promoter activation linked to the cellular identify of OSNs (A) The heatmaps display the co-expression of *Or62*, *Or64*, *Or65*, and *Or67* in four clusters. Rows represent receptors and columns represent cells. The color of the heatmap represents the normalized expression level of OR genes. (B) The genome tracks display the cluster-specific chromatin accessibility around the promoters of *Or62*, *Or64*, *Or65*, and *Or67* in snATAC-seq data, with the promoter region highlighted in light green. The cartoon model depicts the schematic of expression patterns of four gene combinations in clusters 1 – 4. Triangles indicate the transcription start site (TSS) region of *Or62*, *Or64*, *Or65* or *Or67*. The wavy lines represent transcribed RNA molecules, and the solid circles represent the 5’ cap. In gene models, black boxes and thin solid lines represent exons and introns, respectively. The position and relative size of exons and introns were adopted from *A. mellifera* genome annotation and drawn to scale. The positions of the arrows represent the TSS, while the direction of the arrows indicates the direction of transcription. (C) The UMAP plot illustrates the distribution of clusters 1 – 4, employing distinct colors to depict cells belonging to specific clusters. (D-E) (D) Density distribution of transcriptomic Euclidean distances without OR genes (computed on the first 20 PCs) between pairs of OSNs in the same cluster (intra), different clusters (inter), and the same clusters after permutation of all cluster identities prior to distance calculation (intra perm, see methods). Horizontal bars correspond to mean values and dots correspond to median values. (E) Range of Cohen’s d values calculated between the distribution of intra or inter OSNs population pairwise transcriptomic Euclidean distances and each of the distributions of distances after permutation of cluster identities. The red horizontal dashed line corresponds to the Cohen’s d value computed between the distributions of intra and inter OSN population pairwise transcriptomic Euclidean distances. ***p < 0.001. Two-sample Kolmogorov–Smirnov test. (F-G) The genome tracks show the chromatin accessibility around the Netrin receptor UNC5B (F) and zinc finger SWIM domain-containing protein 8 (G) loci. The tracks from top to bottom represents normalized chromatin accessibility around the genes in cluster 1 – 4, gene annotation, peaks, and peak-to-gene links. The vertical bars spanning both panels highlight selected peaks linked to Netrin receptor UNC5B (F, light green: promoter) and zinc finger SWIM domain-containing protein 8 (G, light blue: enhancer) expression that were identified in clusters 1 – 4. Next to the genome track is a violin plot showing gene expression in clusters 1 – 4. (H-K) (H, G) Density distribution of transcriptomic Euclidean distances (computed on the first 10 PCs) within cluster 1 or cluster 2 between pairs of OSNs expressing the same OR gene (same) or different OR genes (diff). (I, K) Range of Cohen’s d values calculated between the distribution of same or different OSNs population pairwise transcriptomic Euclidean distances and each of the distributions of distances after permutation of receptor identities. ns *p* > 0.05. Two-sample Kolmogorov–Smirnov test.

Interestingly, OSNs exhibiting combinatorial expression of these four OR genes were clustered into four classes in our ISAC algorithm, indicating their differential cellular identity (**Figure 5C**). To verify the robustness of transcriptome divergence between the four classes, we compared pairwise transcriptome distances of OSNs within or between different classes (excluding OR genes). Our analysis demonstrated that OSNs within the same cluster (intra) exhibited significantly higher transcriptional similarity compared to those in different clusters (inter) (**Figure 5D-E**). Additionally, detailed analysis revealed differential expression of genes between the four clusters, which was supported by alterations in chromatin accessibility at promoter or enhancer regions (**Figure 5F-G and Figure S5D-F**). In contrast, OSNs in the same cluster, with the same accessible promoter, regardless of expression of the same or different OR genes, demonstrated no significant transcription difference (**Figure 5H-K**). These findings collectively suggest that the cellular identity of OSNs is primarily influenced by the specificity of promoter accessibility, rather than the expressed OR genes. We speculate that the loss of transcription termination sites during OR duplication might be the cause of these co-expressions of multiple OR genes. However, previous studies in insects suggest that the downstream transcripts are byproducts that may not be able to produce functional proteins, which explains their lack of contribution to the cellular identity of OSNs ^46^. Taken together, the polycistronic transcription of adjacent OR genes explains the majority of the co-expression events, supporting the preference for a single functional receptor in *A. mellifera* OSNs ^42,43,47,48^.

### Genomics features and regulatory mechanisms of OR gene expression in honey OSNs

The comprehensive profile of *A. mellifera* OSNs presents an expected and surprising expression pattern of OR genes. In summary, we observed that many OR genes are co-expressed at the RNA level; however, only five out of the 59 OSN classes have the potential to produce multiple proteins (**Figure S6A**). Overall, the OR expression pattern in *A. mellifera* OSNs follows the same rules as *D. melanogaster*, with the majority of neurons expressing a single functional ligand-specific receptor and a small proportion of neurons expressing multiple duplicated OR genes with limited sequence divergence (**Figure 6A**). In *D. melanogaster*, three out of the 34 OSN classes in the adult antenna express multiple OR genes ^16^, as validated at protein level ^43,49–51^. Despite the higher density of OR genes in the *A. mellifera* genome compared to flies, the proportion of functional co-expressed OR genes is surprisingly lower than expected. When considering their duplication time (estimated by sequence distance), the results are even lower than expected. (**Figure S6B**). Based on observations in flies and the evolution trajectory of gene duplication ^51^, we expected to observe a preference for co-expression in newly duplicated OR genes. However, we observed that functional co-expression of OR genes was not enriched or higher in the newly duplicated OR pairs. In contrast, the nonfunctional co-expression of OR genes through polycistronic transcription were highly enriched in the newly duplicated OR genes. Our analysis revealed dramatic differences in the genomic features of single-expression OR genes versus co-expression of OR genes at RNA levels (**Figure 6B-G and Figure S6C-D**), in terms of genome location, sequence similarity, promoter similarity and transcription termination site density. These results indicate, the polycistronic transcription might be operated by these cis-elements. However, there is no significant difference in these features between OR genes that co-express with a single promoter or those with multiple promoters. Therefore, the difference between them might be due to other unknown trans-factors, as there are no genetic features that can distinguish OR genes that co-express with a single promoter from those with multiple promoters. Previous study of *Ir75a*-*c* in flies, and recent single-cell transcriptome analysis in ant OSNs, both suggested that the polycistronic transcription may be a mechanism for repressing the functional expression of downstream OR genes in a tandem duplication array. Our multi-omics profiling demonstrated that the transcription readthrough and co-accessibility of promoters seem exclusive in individual OSNs, indicating the transcription readthrough might repress the downstream genes by remodeling the chromatin accessibility of their promoters. Whether and how polycistronic transcription contributes to OR genes specificity requires further investigation.

**Figure 6.**
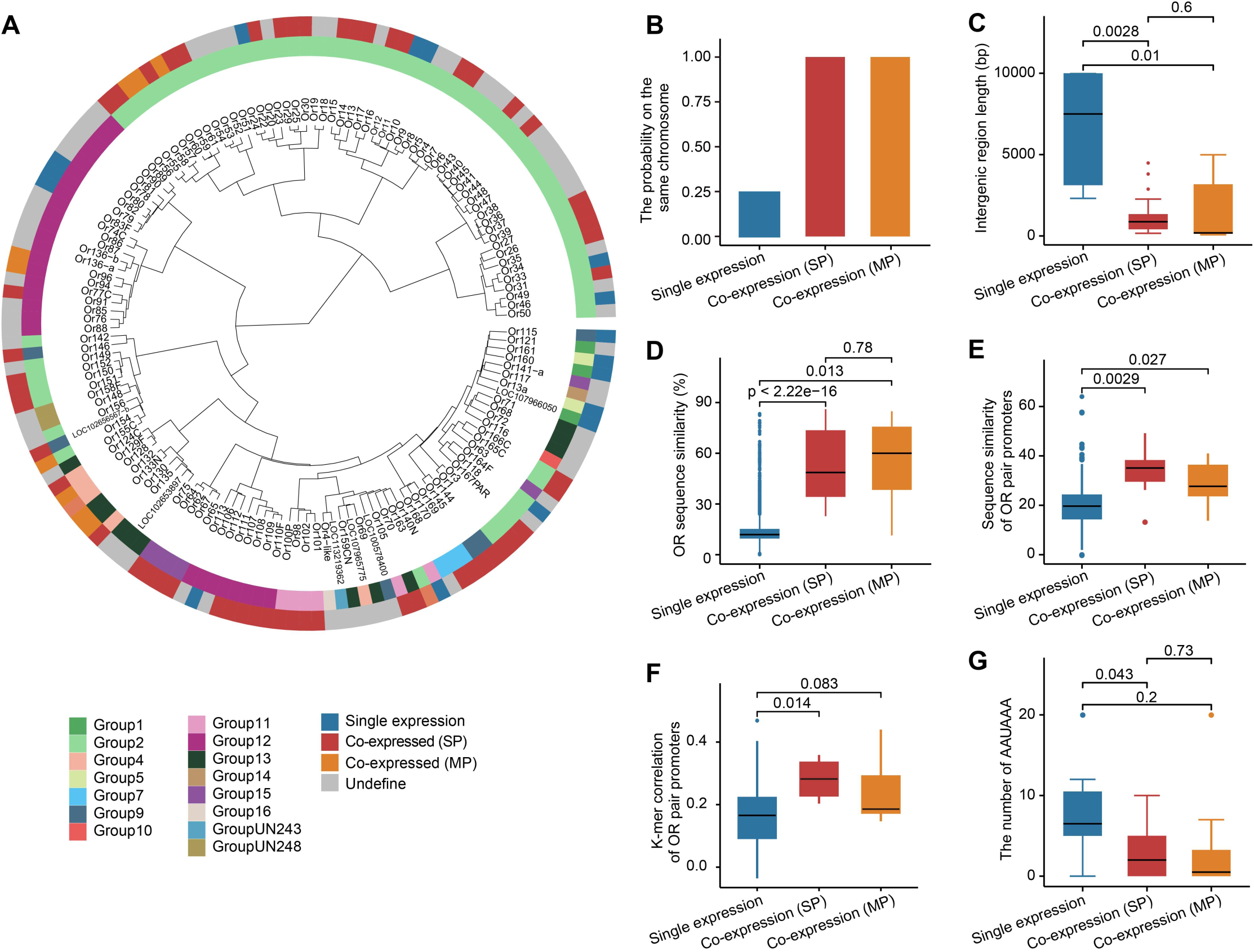
Genomics features associated with the expression pattern of OR genes in *A. mellifera* OSNs (A) The innermost layer of the diagram depicts the phylogenetic relationships among OR genes. Moving outward, the first circle represents the chromosome on which each OR gene is located, and the second circle represents the pattern of gene expression. (B-D) The bar plots illustrate the characteristics of three different expression patterns of OR genes, including the frequency on the same chromosome (B), the length of the intergenic region (C), and the sequence similarity of OR genes (D). The significance of these characteristics was determined using the Wilcoxon Rank Sum test, and the *p*-value is indicated above each plot. Co-expression (SP): OR genes co-expressed with a single promoter; Co-expression (MP): OR genes co-expressed with multiple promoters. (E-F) The box plots represent the sequence similarity (E) and k-mer correlation (F) of OR pair promoters for single expression, Co-expression (SP), and Co-expression (MP) OR genes, compared using the Wilcoxon rank-sum test. The *p*-value is indicated above each plot. (G) The box plot displays the number of AAUAAA sequences in transcription termination region among single expression, Co-expression (SP), and Co-expression (MP) OR genes, compared using Wilcoxon rank-sum test. The *p*-value is indicated above the plot.

## Discussion

In this study, we conducted a systematic profiling of the single-cell transcriptome and chromatin accessibility in antenna tissue from *A. mellifera* workers. Our single-cell multi-omics profiles revealed a comprehensive expression pattern of OR genes in the OSN populations, which were further validated by RNA-FISH. Globally, *A. mellifera* OR genes specialized their expression in OSNs, and their expression specificity also connected to their cellular identities, as demonstrated in *D. melanogaster*. Initially, we observed a higher proportion of OSNs co-expressing multiple OR genes. However, the majority of the co-expression events were polycistronic transcription. Previous studies have demonstrated that the downstream genes in a polycistronic transcript might not be able to be exported to the nucleus and translated into functional proteins ^52^. Therefore, these OSNs might have only one functional receptor, following the one-receptor to one-neuron principles at protein level.

Although only a few insect species have their transcriptome profiles at single cell level for OR expression quantifications, based on current information, especially the new observations in *A. mellifera*, an outgroup species of flies and mosquitos, suggest that the singular receptor per neuron in the olfactory system is the ancestral and normative form in insects. The non-canonical patterns observed in mosquitoes are the minority, but confer a selective advantage under specific conditions ^17^.

The unexpected observation in our study is the functional co-expression of two or three different ligand-specific OR genes, which exhibit multiple co-accessible promoter regions, which are relatively rare in *A. mellifera* OSN classes (only 5 out of 59 clusters in *A. mellifera*). The proportion is even lower than the observations in flies (3 out of 34 clusters) ^16^. These observations refute the previous hypothesis that the co-expression of multiple functional OR receptors might be a temporal state of newly duplicated OR genes that have not yet specialized their expression during the early evolution stage. In addition, these observations also suggest additional mechanisms might be involved in facilitating the divergent expression of newly duplicated OR genes.

Our analysis demonstrated that the newly duplicated OR genes are actually enriched for polycistronic transcription in *A. mellifera* OSNs. Polycistronic transcription was thought to be a random loss or shortening of transcription termination sites, without functional output. The overrepresentation of polycistronic transcription in the tandem duplication OR blocks raises an alternative hypothesis that it may facilitate the OR gene expression specificity by inhibiting the downstream OR genes ^53^. The biochemical mechanism of transcriptional interference is not fully understood but is thought to result from RNA polymerase II complexes transcribing an upstream gene prevent assembly of the transcription machinery at the promoter of a downstream gene. Another mechanism to explain the transcriptional interference is competition during promoter selection ^48,54–56^.

In an evolutionary context, the existence of cis-mediated transcriptional repression in recently duplicated chemosensory genes may provide a substrate for natural selection to favor rapid acquisition of distinct expression patterns (or otherwise pseudogenization of one duplicate). Given the widespread occurrence of chemosensory receptor gene families in arrays in the genomes of most or all animals ^57^, our discoveries may have broad implications for the evolution of their distinct expression patterns. The study of the regulatory interactions between arrays of olfactory receptor genes may reveal general insights into how other types of proteins encoded by tandemly duplicated genes evolve distinct expression patterns.

Our study also demonstrates the power of the single cell multi-omics approach in studying heterogeneous cell populations, specifically for non-model species, where basic genetic approaches are limited or lacking. Integrating information from multiple levels can not only provide efficient features for distinguishing transcriptions via different mechanisms, providing functional inference, but also decipher the mechanisms underlying the gene regulations. Together, our systematic analyses underscore that single-cell multi-omics profiling can systematically deepen our understanding.

## Acknowledgments

We thank Weiwei Zhai (Institute of zoology, Chinese academy of sciences), and Rui Zhang (Sun Yat-sen University) for constructive comments and suggestions on the manuscript, as well as the members of Xu lab for their helpful advice. This work was supported by National Key R&D Program of China (2021YFA1102100 and 2021YFA1102500 to J.X.), National Natural Science Foundation of China (32070644, 32293190 and 32293191 to J.X.), Guangdong Basic and Applied Basic Research Foundation (2022A1515110223 to W.X.ZH.), Fundamental Research Funds for the Central Universities, Sun Yat-sen University (22qntd2621 to W.X.ZH.).

## Author contributions

J.X. conceptualized the project. J.X., W.X.ZH., Y.G.N. and Y.H.L. developed the methodology. J.X., W.X.ZH., Y.G.N. and Y.H.L. conducted the investigation. W.X.ZH., Y.C.X., T.X., P.Y.S., F.L., H.X.ZH. and Q.M. performed visualization. J.X. and W.X.ZH. acquired funding. J.X. administered and supervised the project. J.X. and W.X.ZH. wrote the original draft. W.X.ZH., Y.G.N., Y.H.L., Y.C.X., T.X., P.Y.S., F.L., H.X.ZH., Q.M. and J.X. reviewed and edited the manuscript.

## Materials and methods

### *Apis mellifera* workers collection

*Apis mellifera* workers were obtained from three robust colonies provided by the Institute of Zoology, Guangdong Academy of Sciences (Guangzhou, Guangdong, China). Each colony was housed in a standard Langstroth hive, containing a single hive box with a population of 30,000–40,000 bees, inclusive of one naturally mated queen. To obtain newly emerged workers, capped brood frames were closely monitored until workers emerged from their cells, typically within 1 – 2 hours. In order to gather age-matched samples of nurse bees (9 days post-eclosion) and forager bees (21 days post-eclosion), the adult bees that had freshly emerged from their cells were distinctly marked on the thorax using non-toxic paint and subsequently reintroduced to the hive. On the 9st day post-eclosion, a subset of workers displaying nursing behaviour were collected, identified by their heads being in cells containing larvae. Similarly, on the 21st day post-eclosion, another subset of bees demonstrating foraging behaviour were collected, identified by their return to the hive carrying pollen loads ^1,2^.

### Antennal dissection and single-cell multi-omics library preparation

*A. mellifera* workers were anesthetized on wet ice for a duration of 10 minutes. Subsequently, the antennae were carefully excised using forceps. Prior to dissection, three sterile dishes were prepared: the first containing 100% ethanol, and the second and third filled with phosphate-buffered saline (PBS). The antennae were initially rinsed in the ethanol dish for 5 seconds, followed by a rinse in the PBS dishes for an additional 5 seconds each. The dissection of the antennae was performed under a dissecting microscope using fine dissection scissors, and was conducted in a cold MACS^®^ Tissue Storage Solution (Miltenyi) to ensure tissue preservation. The dissected antennae were then transferred to a pre-chilled DNA LoBind 1.5 mL tube (Eppendorf 022,431,021) containing cold MACS^®^ Tissue Storage Solution. The dissection process for each sample was strictly limited to 120 minutes to maintain the integrity of the nuclei. Post-dissection, the samples were spun down using a benchtop centrifuge, and the MACS^®^ Tissue Storage Solution was carefully removed. The samples were then flash-frozen in liquid nitrogen and stored at −80 °C until further processing for nuclei extraction was required.

Upon thawing on wet ice, the dissected antennae were processed to prepare single-nucleus suspensions, following the protocol outlined in the Fly Cell Atlas study with minor modifications ^3^. The specific steps included centrifugation of the thawed antennae in a benchtop microcentrifuge for 5 – 10 seconds, followed by transferring 1 ml of the homogenized sample into a 1 ml autoclaved dounce (Wheaton #357538). The nuclei were then released by 40 strokes of a loose dounce pestle and 35 strokes of a tight dounce pestle, ensuring that bubble formation was avoided. The sample was then filtered using a 10 µm Flowmi (BelArt #H13680-0040) into a new 1.5 ml Eppendorf tube, which was kept on ice. The filtered sample was centrifuged for 10 minutes at 600 g at 4 °C, and the supernatant was discarded to leave the nuclei. The nuclei were then re-suspended in a desired volume of 1 x Nuclei Buffer supplemented with 1U/µL RNase inhibitor and 1mM DTT. The suspension was mixed by pipetting 20 times to ensure complete re-suspension of the nuclei, and the tube was kept on ice until further processing.

The preparation of the single cell multiome library was conducted in accordance with the guidelines provided in the 10x Genomics Single Cell Multiome ATAC + Gene Expression Reagent Kit user guide. The generation of the dual index libraries was carried out strictly adhering to the standard protocols recommended by 10x Genomics. After the library preparation, the quality and quantity of the libraries were assessed using a Nanodrop spectrophotometer (Thermo Scientific, Waltham, Massachusetts, USA) and an Agilent Bioanalyzer (Agilent Technologies, Santa Clara, California, USA).

### Preparation and sequencing of bulk RNA-seq libraries

Workers at four distinct developmental stages (Newly emerged, Nurse, Forager, and Defender) were gathered from each colony based on their characteristic behaviors as described in previous studies ^4^. Newly Emerged workers were obtained within 1 – 2 hours post-emergence from their cells. Nurses were identified and captured as they moved toward the larval cells. Foragers carrying pollen loads were collected upon their return to the hive. Defenders were identified and captured as they stood at the colony entrance, interacting with incoming bees using their antennae. The antennae from 60 workers representing each labor division were surgically dissected, pooled, flash-frozen in liquid nitrogen, and then stored at −80 °C for subsequent processing.

The frozen antennal tissues were homogenized in TRIzol reagent, and total RNA was extracted following the manufacturer’s instructions (Invitrogen, Life Sciences). The quantity and quality of the extracted RNA were evaluated using a NanoDrop spectrophotometer (Thermo Fisher), a Qubit 4.0 fluorometer (Invitrogen, Carlsbad, CA, USA), and a Bioanalyzer 2100 system (Agilent). All samples demonstrated RNA Integrity Numbers (RINs) exceeding 7, indicating high RNA quality. Subsequently, cDNA libraries were constructed from 1 μg of total RNA from each sample following the standard protocol, and the libraries were quantified using qPCR. The prepared libraries were then sequenced on the DNBSEQ platform (DNBSEQ T7) for 100 cycles from each end of the fragments, resulting in an average of 125 million 100 bp paired-end reads per library.

### Preparation and sequencing of bulk ATAC-seq libraries

The dissection of worker antennae and extraction of nuclei were performed following the methods described in the previous sections. The construction of the bulk ATAC-seq library was carried out according to the methods detailed in previous studies ^3^. In brief, a total of 30,000 cells were subjected to tagmentation using a TruePrepTM DNA Library Prep Kit V2 for Illumina (Vazyme Biotech, TD501) at 37 °C for a duration of 30 minutes. Following tagmentation, the reactions were purified using Zymo DNA Clean & ConcentratorTM 5 columns (Zymoresearch, D4013). The purified samples were then amplified via PCR using a PCR enzyme and index primers (TD202) obtained from Vazyme Biotech. Post-amplification, the libraries were further purified using AMPure XP beads. The purified libraries were then subsequently sequenced on an Illumina Noveseq 6000 platform, generating 150-bp paired-end reads. The quality control of the libraries and the sequencing services were facilitated by Novogene Bioinformatics Technology Co., Ltd.

### Preparation of worker antennae nuclear RNA and RT-PCR

Nuclear-enriched RNA was prepared from worker antennae nuclei following the methods described in the previous sections. The extraction of nuclear RNA was carried out by adding 1 ml of Trizol, following the manufacturer’s instructions. The extracted RNA was then treated with DNase RQ (Promega) using 0.5-1 units of DNase per μg of RNA, as per the manufacturer’s guidelines. The RNA was subsequently recovered through ethanol precipitation, according to standard protocols. RNA concentrations were measured using a Qbit4 (Invitrogen), and 1 μg of RNA was used for cDNA synthesis. The cDNA synthesis was performed using Maxima H Minus reverse transcriptase (Vazyme) and random primers, according to the manufacturer’s protocol. Minus-RT controls were included and analyzed alongside the other samples to ensure efficient DNase treatment. PCR was performed using Phusion (Takara), following the manufacturer’s recommendations. Primers are shown in supplemental table 3.

### Refinement of chemosensory receptors in GTF reference file

Given the rapid expansion and high sequence similarity of the chemoreceptor multi-gene family, their assembly and accurate annotation present significant challenges ^5^. We curated genes in chemoreceptor multi-gene families using the Amel_HAv3.1 genome and annotation from the National Center for Biotechnology Information (NCBI). Initially, we sequenced the antennae of newly emerged workers, nurses, and foragers. Subsequently, we employed IGV-scRNA (https://gitee.com/CJchen/IGV-sRNA) to manually correct the location and structure of published Amel_HAv3.1 odorant receptors (ORs), gustatory receptors (GRs), and ionotropic receptors (IRs) using high mapping quality bulk RNA-seq data. In instances where two genes were erroneously combined into one (e.g., when the distribution of reads on both sides of the BAM file in bulk RNAseq appeared to be very different), we used the suffixes "-a" and "-b" to distinguish them.

### Identification of OR, GR, and IR genes via domain scanning of protein sequences

During the gene correction process, some genes were split into multiple genes. Pfamscan ^6,7^ was utilized to identify protein sequences containing the odorant receptors domain (7tm_6), gustatory receptors domain (7tm_7 and Trehalose_recp), and ionotropic receptors domain (ionotropic receptor) ^8^. We identified a total of 150 OR genes, 16 GR genes, and 19 IR genes in the Amel_HAv3.1. Following this, we aligned the sequences of the newly identified chemosensory receptors, particularly the 150 OR genes, with previously documented *A. mellifera* OR gene sequences (supplemental table 1). In cases where the sequence identity (pident) exceeded 90%, we adopted the nomenclature from the referenced study ^9^ for the respective OR genes. Otherwise, we retained the current NCBI nomenclature.

### Generation of gene and peak matrix from sequencing data

The ‘cellranger-arc mkref’ tool (version 2.0.0) from 10x Genomics was utilized to create a reference package using the Amel_HAv3.1 genome and the modified annotation GTF file. The raw sequencing scRNA-seq and scATAC-seq FASTQ files were aligned to the Amel_HAv3.1 reference genome using ‘cellranger-arc count’, and the barcoded count matrices of gene expression and chromatin accessibility data were generated. All ambiguous Unique Molecular Identifiers (UMIs) that mapped to multiple genes were removed from the BAM file using custom scripts with samtools and pysam to prevent UMI double-counting. The gene expression matrix was then recalculated by counting the number of distinct UMIs per cell barcode for each gene ^8^.

### Decontamination of ambient RNA in *A. mellifera* antennae

The majority of the cells in the *A. mellifera* antennae are olfactory sensory neurons (OSNs). As the marker gene (odorant receptor co-receptor, *Or2*) of OSNs is widely expressed, it can easily become ambient RNA and be amplified along with a cell’s native mRNA. To estimate and remove the percentage of cross-contamination within each cell due to ambient RNA or other experimental factors, we used ‘decontX’ from the ‘celda’ R package (version 1.10.0) ^10^. The decontaminated reads were rounded using the ‘base:round’ function in R, and the decontaminated matrices were generated using the DropletUtils package (version 1.10.3) ^11^.

### Preprocessing and quality control of scRNA- and scATAC-seq data

Decontaminated or raw count matrices and ATAC matrices for each sample were loaded into Signac ^12^. DoubletFinder (version 2.0.3) was employed to identify and remove doublets ^13^, with the pK that yielded the maximum AUC. Cells were filtered based on criteria of 300-30000 RNA transcripts, 300-100000 ATAC fragments, less than 5% mitochondrial reads, less than 2 nucleosome signals, and TSS enrichment greater than 1. Peak calling was performed using macs2 (version 2.2.5) ^14^. In total, 8,868 antennal cells that met the selection criteria were identified.

### Integrative analysis of single-cell multi-omics identifies neurons in worker antennae

Initially, rPCA was used to identify integration anchors using the RNA modality. The same set of anchors was then used to correct the cell embeddings for both the RNA data (by running IntegrateData, followed by ScaleData and RunPCA on the integrated assay), and for the ATAC data (by running IntegrateEmbeddings with ‘lsi’ reductions). Subsequently, FindMultiModalNeighbors was run using the integrated RNA data and the corrected LSI as inputs, which were integrated dimension reductions used for multimodal clustering. This approach identified 34 clusters containing 7,332 neurons based on the chromatin accessibility in promoter regions and the expression levels of known neuronal marker genes (e.g., synaptotagmin 1 (*syt1*), embryonic lethal abnormal vision (*elav*), cadherin-N (*cadN*), and bruchpilot (*brp*)).

### Identification of OSNs from neurons

In the decontaminated matrix, the ambient RNA from the OSN co-receptor was removed. This led to the identification of 7,212 OSNs exhibiting high chromatin accessibility in promoter regions and expression levels of *Or2* (orthologous to the *D. melanogaster Orco*). As depicted in Figure 1I, the proportion of OSNs in the antennae of workers significantly higher than that in *D. melanogaster* ^3^ and *Ae. Aegypti* ^15^.

### Measurement of transcriptome distance

In line with previous studies on olfactory neurons in mice (16), we defined the transcriptome distance as the cosine distance (1-cosine similarity) between the principal component scores (top 50 PCs) of OSN pairs to measure cell similarity based on gene expression. The cosine distance ranges from 0 (indicating similar transcriptomes at a 0-degree angle) to 1 (indicating orthogonality at a 90-degree angle) and up to 2 (indicating opposing transcriptomes at a 180-degree angle).

### Identification of OSN subtypes using iterative subcluster after clustering

Due to the high sparsity of ATAC data, the OSN clusters were re-integrated and re-clustered by the RNA data with the top 500 highly variable genes from sctransform. The validity of cluster separations was evaluated using a dotplot for chemosensory receptor gene expression and a heatmap for global transcript distance similarity. This evaluation essentially determined whether a cluster separation identified populations with distinct expression of chemosensory genes. The high similarity between OSN clusters necessitates constant iterative subclustering for distinction. To enhance efficiency, we clustered and built trees for subclustering groups based on the transcriptome distance between them, and divided them into four cell groups for further clustering. For each cell group, an appropriate number of variable features was selected for further PCA dimension reduction and clustering. In cases where cells expressing different OR genes were clustered together, additional clustering was performed. Subsequently, transcriptome distances within and between the sub-clusters were analyzed to determine potential over-clustering. A total of 115 populations of OSNs were initially identified. A stringent criterion was then applied, retaining only 59 robust OSN populations characterized by a cell count exceeding 30 and exhibiting significant chemoreceptor gene expression (defined as pct.exp > 40% and avg.exp.scaled ≥ 2.5).

### Identification of the OR gene expressed in each OSN

The dot plot of cluster-enriched chemosensory receptors was created using the DotPlot function in Seurat. The scaled mean normalized expression and expression percentage of each chemosensory receptor was calculated by the DotPlot function. For chemosensory receptors, we define ‘expression’ of a gene within a cluster by pct.exp > 40 and avg.exp.scaled ≥ 2.5 in dotplot data. If a cluster exhibits more than one chemosensory receptor meeting the specified criteria, it is categorized as a multiple-OR-cluster. In our study, there were a total of 28 co-expressed OSN clusters, accounting for approximately 47% of the total, a proportion comparable to the 45% co-expression ratio observed in *Ae. Aegypti* ^15^, and significantly higher than the 11% observed in *D. melanogaster* ^16^.

### Mean ATAC signal in the OR promoter regions within co-expression clusters

First, we extracted the multiple-OR-clusters based on Figure 2B. If there were single-OR-clusters expressing the same OR genes as those in the extracted multiple-OR-clusters, they were also extracted. Then, we obtained the mean of reads for each cluster from the ATAC matrix in the promoter region of the expressed OR genes and remove clusters with weak ATAC signals and their corresponding expressed genes.

### ATAC profiling of co-expressed olfactory sensory neurons

For clusters with multiple active promoters, such as the expression of *Or154* and *Or163* in cluster 36, we defined three cell groups were defined: cells exclusively expressing *Or154*, cells exclusively expressing *Or163*, and cells co-expressing both.

The subset-bam (v1.1.0) (https://github.com/10XGenomics/subset-bam) tool was employed to extract barcode-specific BAM files, which were subsequently converted to BEDGraph using genomecov of bedtools (v2.26.0) ^17^. Normalization and sorting of BEDGraph files were performed with a custom script, followed by conversion to BigWig format using bedGraphToBigWig. A track plot illustrating ATAC accessibility and RNA expression levels within the 500 base upstream of *Or154* to 500 base downstream of *Or163* was generated using ggplot2 ^18^. For a cluster with only one promoter, such as cluster 24 expressing *Or25*, *Or26* and *Or27*, the cells co-expressing *Or25*, *Or26* and *Or27* were extracted, and the bigwig file was extracted and plotted using the same method as described above.

### Processing of TF motifs

Transcription factor (TF) motifs were obtained from the CIS-BP V2.00 Database ^19^, Apis_mellifera Archive. CIS-BP motifs were associated with TF gene symbols by using bioDBnet (https://biodbnet-abcc.ncifcrf.gov/) to link the provided DBIDs to the gene symbols. This resulted in the acquisition of 640 motifs from Apis_mellifera Archive dataset. Given the presence of redundant motifs in the CIS-BP V2.00 database, we subsequently refined the TF motifs to one motif per gene, ultimately yielding 461 distinct motifs.

### TF candidate screening

Due to the sparsity of single-cell data, we aimed to obtain the most comprehensive promoter position information by combining the peaks derived from both single-cell and bulk ATAC-seq data. To identify the TF candidates, we utilized ChIPseeker for peak annotation to the nearest genes ^20^. Upon extracting the peaks annotated to the OR genes, we filtered based on the distance from the transcription start site (TSS), assuming that peaks within 650 base pairs indicate promoter regions.

Subsequently, the corresponding sequences were obtained based on the position of the promoter peak. These sequences were then imported into TOBIAS ^21^ and scanned for enriched TF motifs on each sequence using the TFBScan function. The appropriate candidate TFs were then selected based on Tau score, expression level, logFC value and predicted binding score. The Tau score serves as a statistic to assess the specificity of expression ^22^, and is calculated using the formula:

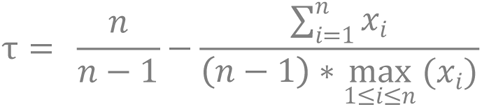

The expression level of TFs was determined from the Seurat RNA assay. The clusters other than the one under consideration were integrated as the ‘Other’ cluster. The logFC value of pairwise comparison between the visited and ‘Other’ cluster was obtained using the FindMarkers function.

The binding score, obtained from the TOBIAS output, represents the score of a TF motif against a genomic sequence, indicating how well the motif fits the sequence. A higher binding score indicates a better match.

### Yeast one-hybrid assay

The yeast one-hybrid (Y1H) system was employed to investigate the transcriptional activation of the *Or128* and *Or129F* promoters by the HTH transcription factor. The open reading frame of HTH was constructed in combination with the pGADT7 vector (Clontech), which served as the effector construct. The promoter of *Or128* and *Or129F*, a 1,000-1,500 bp DNA sequence upstream of the start codon of each gene, was amplified using genomic PCR, and then the PCR products were purified using a PCR kit. The promoter region of each gene was cloned into the pHIS2 vector (three tandem repeats). Positive clones were further confirmed by sequencing. The resulting recombinants were co-transformed into the Y187 yeast strain using an Ex-Yeast Transformation Kit. The yeast colonies were then transferred to plates containing SD/-Trp/-Leu and SD/-His/-Trp/-Leu media, supplemented with 25 mM 3-AT, and allowed to grow at 30°C for 3 days.

### Transcriptomic euclidean distance calculation among clusters

To test whether OSN clusters are transcriptionally dissimilar from each other, we computed the pairwise transcriptomic Euclidean distances between pairs of clusters on the first 20 PCs of clusters 1, 2, 3, and 4, which were devoid of OR gene counts. These distances were sorted into two categories: distances between pairs of OSNs in the same cluster (intra) and distances between pairs in different clusters (inter). To compare and measure the dissimilarity between these two distributions of pairwise transcriptomic Euclidean distances, the Cohen’s d was calculated between the intra and inter distributions using the cohen.d function of the effectsize R package version 0.8.6 with default settings. Moreover, to test if the difference in the distributions of Euclidean distances between pairs of OSNs in the same or different clusters was not due to random effects, we generated a permuted dataset by randomly redistributing the cell identities of the top 20 PCs and recalculated pairwise distances for OSNs with same random identities (same vs perm). By iterating this process, the random distribution was estimated from 1000 permuted datasets. As described above, the Cohen’s d was then calculated between the same (same) or different (diff) OSN population transcriptomic Euclidean distance distributions and each of the permuted distributions. This resulted in 1000 Cohen’s d values for each set of comparisons (i.e., same versus same perm and diff versus same perm). The two distributions of Cohen’s d values were then compared using a two-sample Kolmogorov–Smirnov test. Inaddition, we calculated the distributions of transcriptomic Euclidean distances within cluster 1 and cluster 2 for OSN pairs expressing the same OR gene versus those expressing different OR genes, using a similar approach as described above.

### RNAscope in situ hybridization

*A. mellifera* workers were anesthetized on wet ice, and their antennae were carefully excised using sharp forceps. The antennae were then placed in a 1.5 mL tube and fixed with 4% paraformaldehyde at 4 °C overnight. The specific orientations of each sample, as indicated in the main text and figure legends, were maintained to ensure that sectioning was done along the desired axis. Samples from the same stage were embedded and sectioned together using Tissue-Teck OCT, and then stored at −80 °C until further use. Cryosection was performed with the specified slice thickness using a Leica CM1950 cryostat.

One potential limitation of these experiments is the cross-reactivity of RNA-FISH probes due to the high level of sequence identity across paralogues, which also precluded double RNA-FISH experiments to examine receptor co-expression. To address this, we generated single-base resolution probes. Spatial FISH Ltd (Shenzhen, Guangdong, China) designed the specific probes for target RNA, with all specific primers listed in supplemental table 2. Samples were fixed with 4% paraformaldehyde and enclosed within a reaction chamber for subsequent reactions. Following dehydration and denaturation with methanol, the hybridization buffer containing specific targeting probes was added to the chamber and incubated at 37 °C overnight. Samples were then washed three times with PBST before the target probes were ligated in ligation mix at 25 °C for 3 hours. Subsequently, samples were washed three times with PBST and subjected to rolling circle amplification by Phi29 DNA polymerase at 30 °C overnight. The fluorescent detection probes in hybridization buffer were then applied to the samples. Finally, samples were dehydrated with a series of ethanol washes and mounted in SlowFade Diamond (Thermo Fisher S36972) on glass slides for confocal imaging. All confocal images were captured through a Z-stack scan from the most anterior to the most posterior of the antenna using the OLYMPUS FV31S-SW. Images were then processed with ImageJ and Adobe Illustrator. For quantification, a minimum of three workers (six antennae) were used.

### Compute the length of intergenic regions

The intergenic region for single expression genes is defined as the distance from the transcription termination site of one single expression OR gene to the transcription start site of the next OR gene. For co-expression OR genes with a single accessible promoter, the intergenic region is defined as the distance from the transcription termination site of the first OR gene to the transcription start site of the next OR gene. Similarly, for co-expression OR genes with multiple accessible promoters, the intergenic region is defined as the distance from the transcription termination site of the first OR gene to the transcription start site of the next OR gene.

### Analysis of amino acid sequence similarity

Initially, we extracted the amino acid sequences of significantly expressed OR genes. Multiple sequence alignment was conducted using the muscle function from the R package muscle ^23^. The distances between sequences were then calculated using the maskGaps and stringDist functions from the Biostrings package ^24^. The phylogenetic tree was constructed using the hclust and as.phylo functions. Subsequently, the tree was visualized using the ggtree package ^25^.

### The k-mer analysis in promoter regions of OR gene pairs

Jellyfish ^26^ was employed for tallying the occurrence of k-mers (k=5,7,9,11) within each OR gene’s promoter region. Following this, co-expressed OR gene pairs exhibiting multiple promoters versus those expressing singularly were extracted. The Wilcoxon rank-sum test was used to compute the k-mer correlation between different types of OR gene pairs.

### Compute the length of intergenic regions and count the number of polyadenylation signals (AAUAAA)

The intergenic region for single expression genes is defined as the distance from the transcription termination site of one single expression OR gene to the transcription start site of the next OR gene. For co-expression OR genes with a single accessible promoter, the intergenic region is defined as the distance from the transcription termination site of the first OR gene to the transcription start site of the next OR gene. Similarly, for co-expression OR genes with multiple accessible promoters, the intergenic region is defined as the distance from the transcription termination site of the first OR gene to the transcription start site of the next OR gene. When calculating the number of polyadenylation signals in the transcription termination sequences of OR genes for different expression patterns, the first step is to extract the transcription termination sequences for each OR gene (i.e., the intergenic regions as defined above). Subsequently, these sequences are translated into RNA sequences, and the count of AAUAAA signals is determined.

### Assessment of structural similarity among different OR genes using RMSD

Upon obtaining all OR gene PDB files from Uniprot ^27^, we used Biopython to calculate pairwise Root Mean Square Deviation (RMSD) values between OR genes ^28–30^. This analysis is used to evaluate the structural similarity of proteins, where smaller RMSD values indicate greater similarity, and larger values suggest increased structural divergence.

### Characterize the expression patterns of recently duplicated OR gene pairs across different amino acid distances

We extracted recently duplicated gene pairs from the results of multiple sequence alignment of OR amino acid sequences. For a given pair of aligned sequences, each substitution was scored according to the Miyata amino acid replacement matrix, and insertions were scored as the mean replacement scores of each additional amino acid. The sum of these scores represented the pairwise sequence difference. Sequence differences were then divided into 10 equal bins, and we tallied the number of expression patterns for OR pairs in each bin.

## Data availability

Raw FASTQ files and processed files, including gene and peak matrix, can be downloaded from NCBI GEO under accession number GSE248097 (To review GEO accession GSE248097: Go to https://www.ncbi.nlm.nih.gov/geo/query/acc.cgi?acc=GSE248097 Enter token yvwtmwoujjoprcn into the box).

## Code availability

All analysis codes for generating the figures are available in https://github.com/nieyage/honeybee_ORN.

## Figure legends

**Figure S1.**
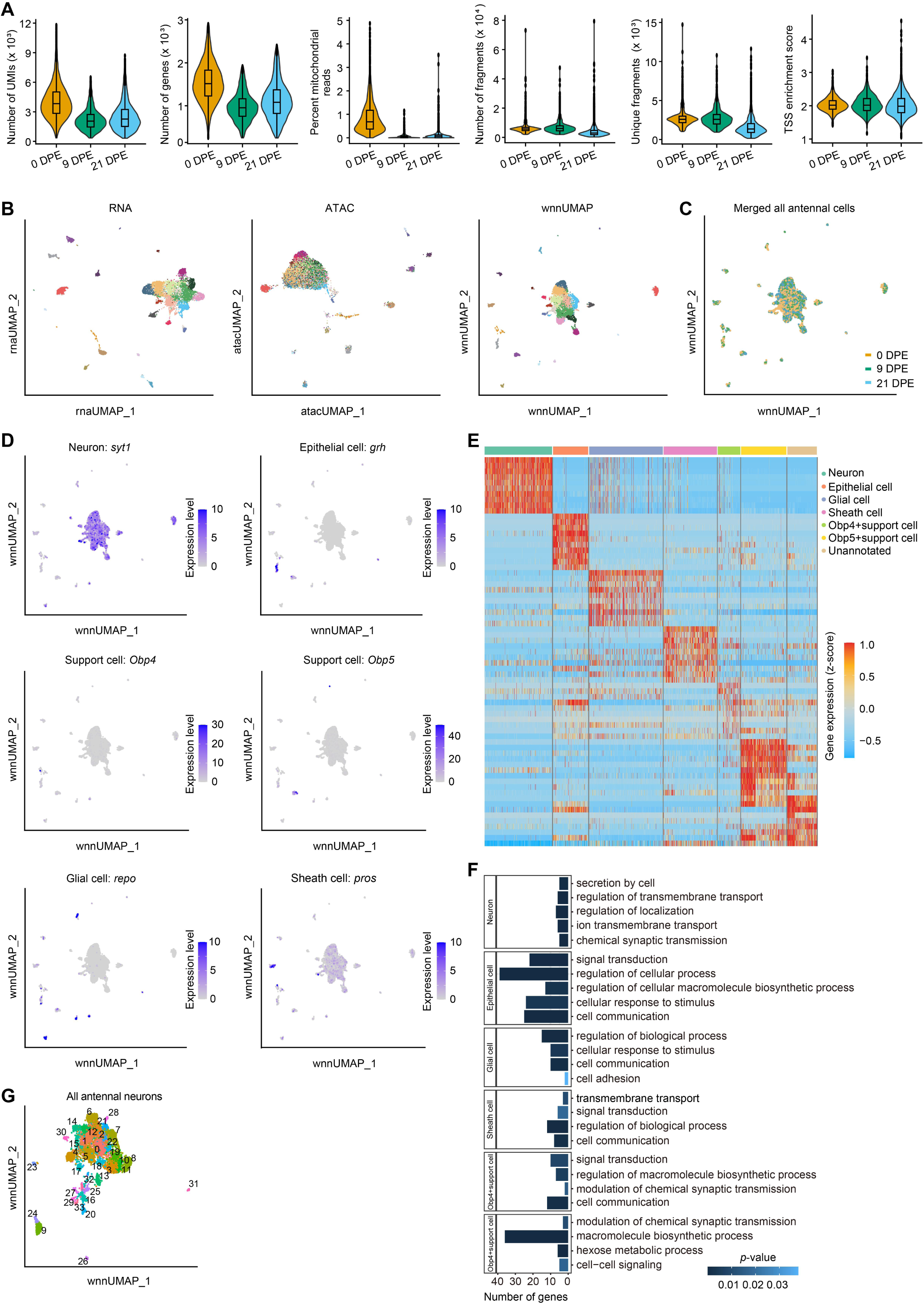
Single-cell multi-omics analysis and quality control (A) Violin plots display the distributions of quality metrics for multi-omics data in each sample, including the number of UMIs, the number of genes, the percentage of mitochondrial reads, the number of fragments, the unique fragments, and the normalized accessibility around transcription start site (TSS) enrichment scores. (B) The UMAP plot represents the clustered results of quality-controlled snRNA, snATAC, and integrated single-cell multi-omics dataset, with different clusters color-coded. (C) The wnnUMAP plot shows the full single-cell multi-omics dataset after quality control, colored by different samples. (D) Expression is mapped to wnnUMAP plots for specific markers, such as *syt1* for neurons, *grh* for epithelial cells, *repo* for glial cells, *pros* for sheath cells, and *Obp*4/5 for Obp4/5+ support cells. (E) The heatmap displays the top cluster-specific genes in seven cell types. Each row represents a cluster-specific gene, and each column represents a cell. The color indicates normalized expression (z-score). (F) Gene Ontology (GO) functional enrichment analysis of cluster-specific genes for each cell type. The top 5 most significantly enriched GO terms are shown, along with the significance of enrichment and the number of genes corresponding to each term. The color of the bar plot represents the *p*-value. (G) wnnUMAP plot represents all antennal neurons annotated by clusters.

**Figure S2.**
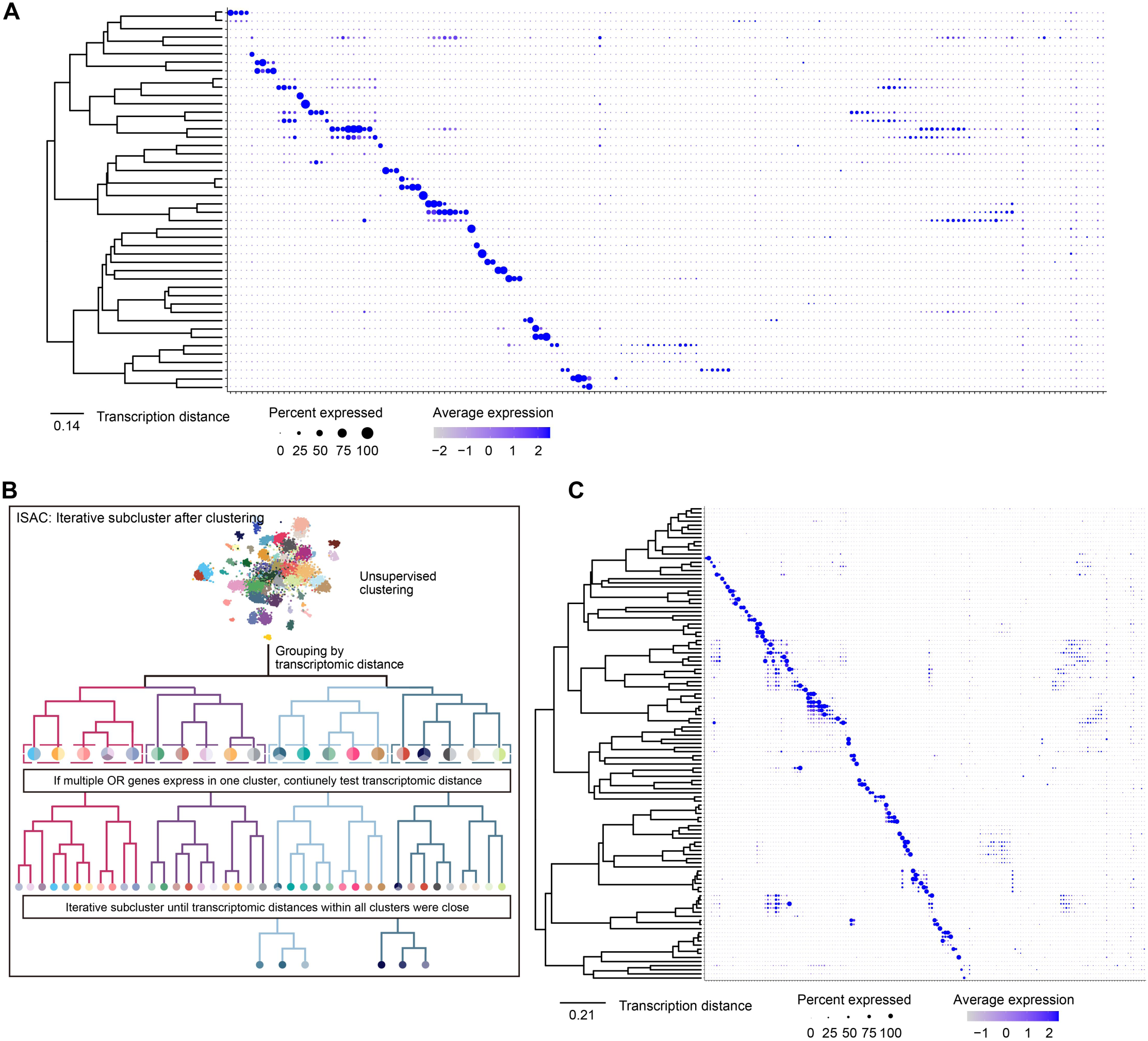
OSNs clustering using unsupervised methods (A) The tree represents the transcriptomic distance between OSN clusters before using Iterative Subcluster After Clustering (ISAC). The dotplot shows the expression of OR genes in each population before ISAC. Clusters are indicated by rows, and OR genes by columns. (B) Schematic of ISAC, an unsupervised algorithm for identifying clusters. (C) The tree represents the transcriptomic distance between OSN clusters after ISAC, with the dotplot showing the expression of OR genes in each population after ISAC. Clusters are indicated by rows, and OR genes by columns.

**Figure S3.**
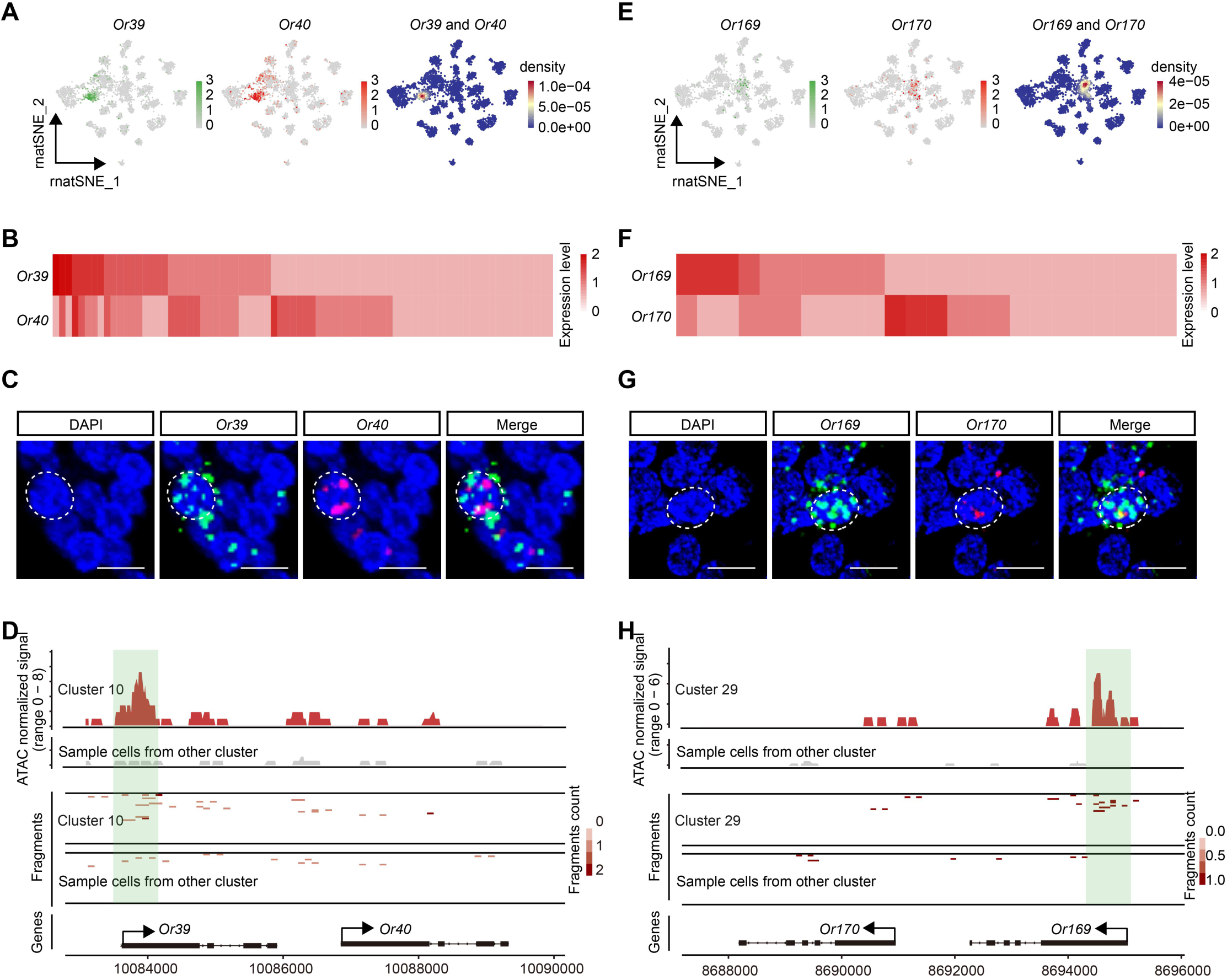
OR genes co-expressed with single active promoter (A-D) *Or39* and *Or*40 are co-expressed in cluster 10. Details are from Figure 3A-D. The order of the panels and legends is the same as in (A-D). (E-H) *Or169* and *Or170* are co-expressed in cluster 29. Details are from Figure 3A-D. The order of the panels and legends is the same as in (A-D).

**Figure S4.**
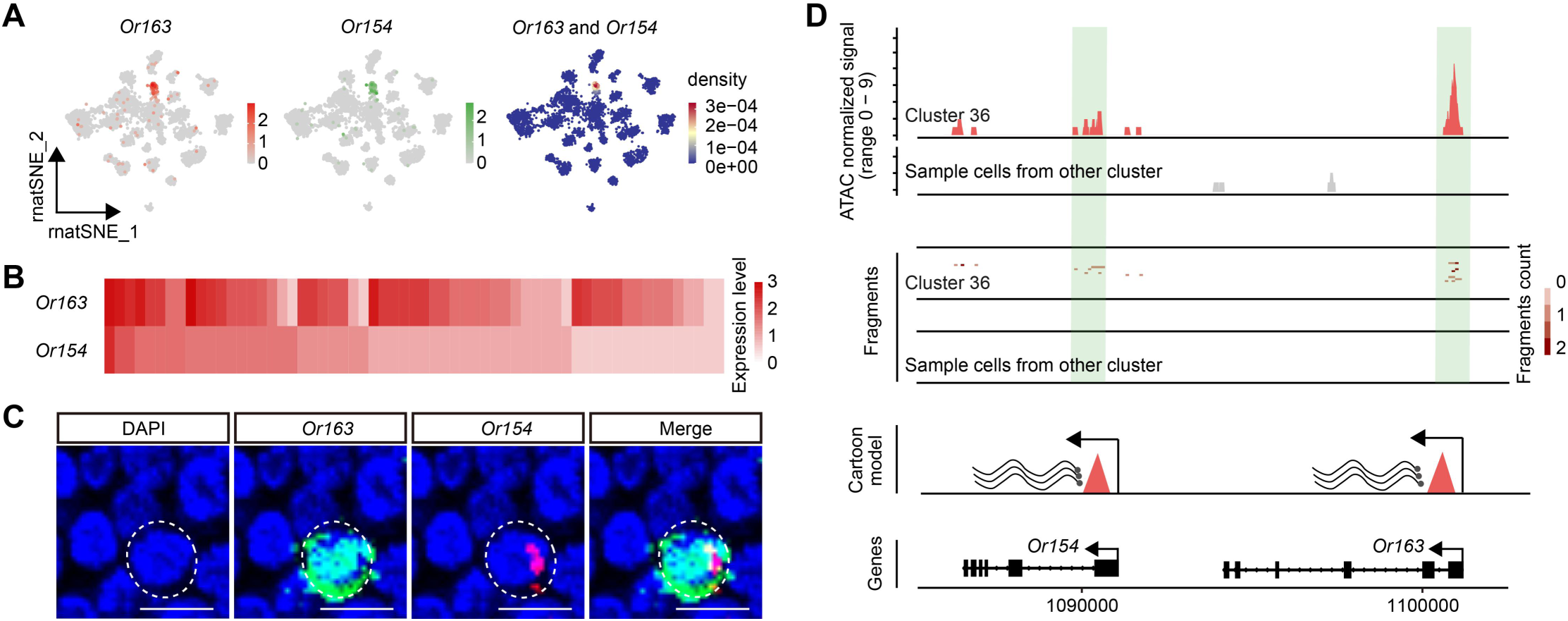
OR genes co-expressed with multiple active promoters (A-D) *Or163* and *Or154* are co-expressed in cluster 36. Details are from Figure 4A-D. The order of the panels and legends is the same as in (A-D).

**Figure S5.**
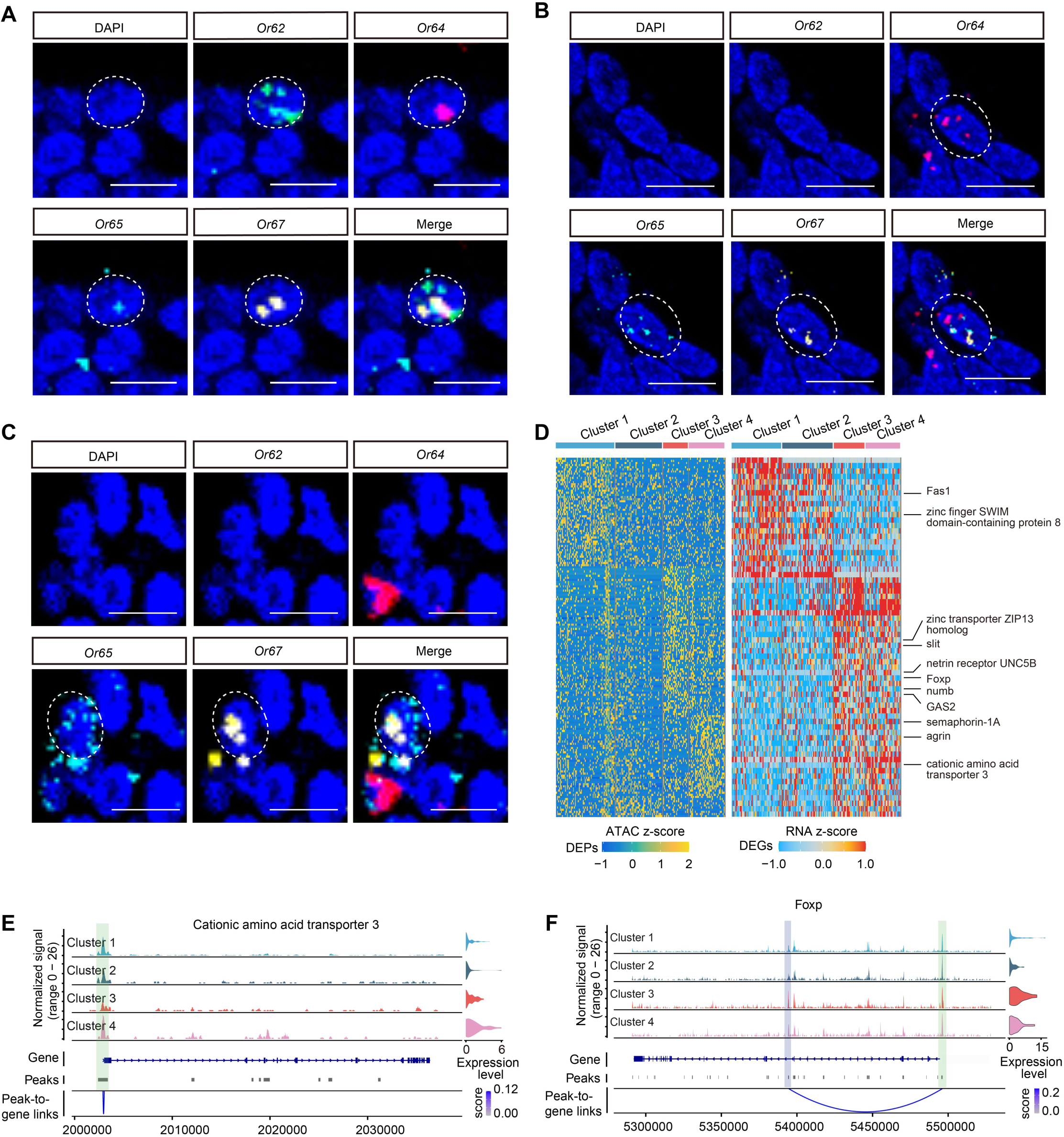
Combination expression of OR gene in a duplication array (A-C) High magnification images of a four-channel RNA-FISH experiment on a transversal section from an antenna of worker incubated with antisense RNA probes for *Or62* (green), *Or64* (red), *Or65* (cyan), and *Or67* (yellow). The section was counterstained with DAPI (blue). The dashed circles represent the nuclei of neurons. Scale bars: 5 µm. (D) (left) The heatmap shows differentially accessible peaks (DEPs) among the four different clusters. The color indicates the z-transformed chromatin accessibility of the DEPs. (right) The heatmap displays the scaled expression of top differentially expressed genes (DEGs) in the four different clusters, respectively. The color indicates the normalized expression. Each row represents a DEP or DEG, and each column represents a cell. Typical axon guidance-related genes are shown on the right of the heatmap. (E-F) The genome tracks show the normalized chromatin accessibility around Cationic amino acid transporter 3 (E), and Foxp (F) loci. The tracks from top to bottom represent the normalized chromatin accessibility around the genes in cluster 1 – 4, gene annotation, peaks, and peak-to-gene links. The vertical bars spanning both panels highlight selected peaks linked to Cationic amino acid transporter 3 (E, light green: promoter) and Foxp (F, light blue: enhancer) expression that were identified in cluster 1 – 4. Beside the genome track is the violin plot showing gene expression in cluster 1 – 4.

**Figure S6.**
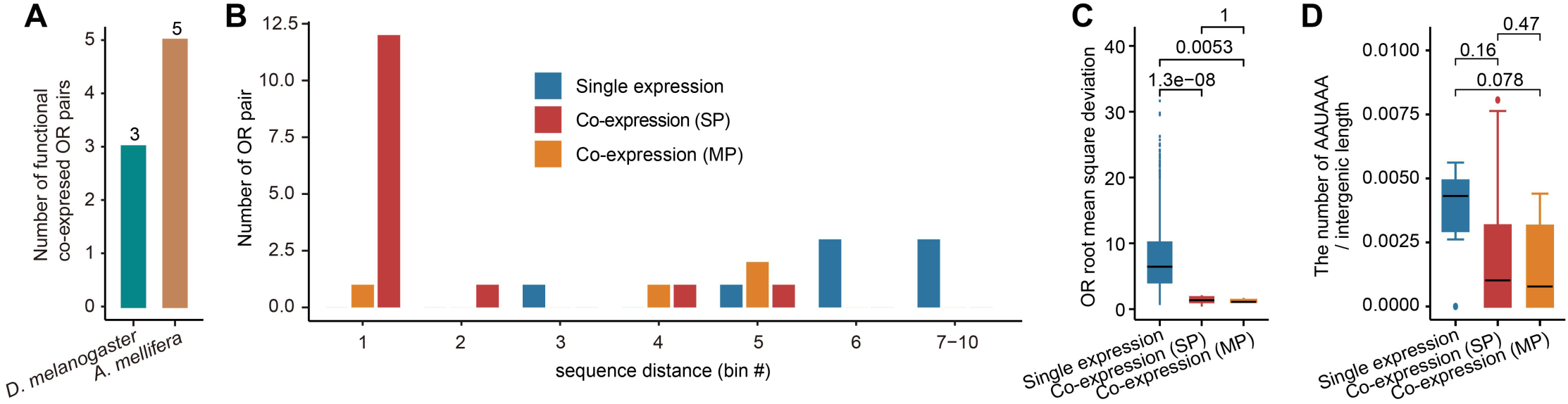
Expression pattern and genomics features of OR gene in *A. mellifera* (A) The bar plot shows the number of functional co-expressed OR gene pairs in *D. melanogaster* and *A. mellifera*. (B) Distribution of number of OR gene pairs per bins of sequence distances, for single expression, Co-expression (SP), Co-expression (MP) OR gene pairs belonging to the close gene cluster in *A. mellifera*. Co-expression (SP): OR genes co-expressed with a single promoter; Co-expression (MP): OR genes co-expressed with multiple promoters. Sequence distances: sum of Miyata amino acid replacement scores between protein sequences. (C) The bar plot illustrate the root mean square deviation (RMSD) of OR gene proteins of three different expression patterns of OR genes. The significance of these characteristics was determined using the Wilcoxon Rank Sum test, and the *p*-value is indicated above the plot. (D) The box plot displays the ratio of AAUAAA sequences in transcription termination region among single expression, Co-expression (SP), and Co-expression (MP) OR genes, compared using Wilcoxon rank-sum test. The *p*-value is indicated above the plot.

