## Supplemental Table 1 for "Singular Odorant Receptor Expression Orchestrated by Promoter Activation Specificity in *Apis Mellifera* Olfactory Sensory Neurons": Supplemental table 1.pdf

Supplemental table 1. Names for OR genes

| OR gene names in NCBI | Robertson, H. M., & Wanner, K. W. (2006) | Pident | OR gene names in this study | Group | Start | End | Width | Strand | Transcript ID |
|---|---|---|---|---|---|---|---|---|---|
| LOC100578400 | NA | 0 | LOC100578400 | Group9 | 9370774 | 9373667 | 2894 | - | XM_026442780.1 |
| LOC102653858 | NA | 0 | LOC102653858 | Group13 | 1003533 | 1013279 | 9747 | + | GSAman00023 |
| LOC102653897 | NA | 0 | LOC102653897 | Group13 | 919759 | 924795 | 5037 | + | XM_026444719.1 |
| LOC102653979 | NA | 0 | LOC102653979 | GroupUN2 | 198771 | 200342 | 1572 | - | XM_026445955.1 |
| LOC102656567-b | NA | 0 | LOC102656567-b | Group9 | 9749864 | 9751731 | 1868 | - | GSAman00027 |
| LOC107965775 | NA | 0 | LOC107965775 | Group4 | 790666 | 794062 | 3397 | - | GSAman00014 |
| LOC107966050 | NA | 0 | LOC107966050 | Group1 | 10753650 | 10756361 | 2712 | - | XM_026443848.1 |
| LOC113219362 | NA | 0 | LOC113219362 | GroupUN2 | 66474 | 69178 | 2705 | - | XM_026445982.1 |
| LOC725205 | AmOr1 | 99.75 | Or1 | Group2 | 9976746 | 9979176 | 2431 | + | XM_026439127.1 |
| Or10 | AmOr10 | 99.494 | Or10 | Group2 | 10004087 | 10005752 | 1666 | + | NM_001242961.1 |
| LOC102656805 | AmOr100P | 100 | Or100P | Group11 | 11132256 | 11134536 | 2281 | - | XM_026443829.1 |
| LOC102655180 | AmOr101 | 99.257 | Or101 | Group11 | 11129371 | 11132005 | 2635 | - | XM_016915293.2 |
| LOC102655147 | AmOr102 | 100 | Or102 | Group11 | 11125399 | 11128016 | 2618 | - | XM_016915299.2 |
| Or105 | NA | 0 | Or105 | Group11 | 11116384 | 11118421 | 2038 | - | GSAman00066 |
| LOC102653810 | AmOr106 | 100 | Or106 | Group12 | 4912295 | 4917424 | 5130 | + | XM_006562209.3 |
| Or107 | AmOr107 | 100 | Or107 | Group12 | 4918290 | 4921395 | 3106 | + | NM_001242987.1 |
| LOC102653716 | AmOr108 | 100 | Or108 | Group12 | 4922227 | 4926859 | 4633 | + | XM_006562208.3 |
| Or109 | AmOr109 | 99.744 | Or109 | Group12 | 4927866 | 4930728 | 2863 | + | NM_001242988.1 |
| Or11 | AmOr11 | 100 | Or11 | Group2 | 10008722 | 10010579 | 1858 | + | NM_001242962.1 |
| LOC102653865 | AmOr110F | 99.743 | Or110F | Group12 | 4931938 | 4934844 | 2907 | + | XM_006562210.3 |
| LOC100577888-a | AmOr111 | 100 | Or111 | Group12 | 4936142 | 4938820 | 2679 | + | GSAman00070 |
| LOC100577888-b | AmOr112 | 99.733 | Or112 | Group12 | 4940729 | 4943724 | 2996 | + | GSAman00077 |
| LOC100577888-c | AmOr113 | 99.486 | Or113 | Group12 | 4945340 | 4948817 | 3478 | + | GSAman00078 |
| Or115 | AmOr115 | 100 | Or115 | Group9 | 11551015 | 11554325 | 3311 | + | NM_001242989.1 |
| LOC100576462 | AmOr116 | 94.915 | Or116 | Group10 | 9480194 | 9482328 | 2135 | + | XM_003250778.4 |
| LOC100576734 | AmOr117 | 100 | Or117 | Group14 | 1435069 | 1437474 | 2406 | + | XM_026444979.1 |
| LOC102654074 | AmOr118 | 100 | Or118 | Group2 | 1056169 | 1059703 | 3535 | + | XM_006572121.2 |
| Or12 | AmOr12 | 100 | Or12 | Group2 | 10013251 | 10014901 | 1651 | + | NM_001242963.1 |
| LOC102656429 | AmOr121 | 91.96 | Or121 | Group1 | 14326374 | 14328764 | 2391 | + | XM_026444854.1 |
| LOC102653782 | AmOr124C | 94.444 | Or124C | Group4 | 6019625 | 6025554 | 5930 | - | XM_026440490.1 |
| LOC102656904 | AmOr128 | 92.288 | Or128 | Group4 | 6016570 | 6019462 | 2893 | - | XM_006561964.3 |
| LOC102656221 | AmOr129F | 92.288 | Or129F | Group4 | 6013575 | 6016404 | 2830 | - | XM_026440491.1 |
| Or13 | AmOr13 | 99.467 | Or13 | Group2 | 10015224 | 10017204 | 1981 | + | XM_026446349.1 |
| Or130 | AmOr130 | 92.602 | Or130 | Group4 | 6010255 | 6013388 | 3134 | - | XM_016911570.2 |
| LOC102653615 | AmOr132 | 100 | Or132 | Group13 | 935464 | 940440 | 4977 | + | XM_026444646.1 |
| LOC102653695 | AmOr133N | 100 | Or133N | Group13 | 943330 | 948576 | 5247 | + | XM_026444679.1 |
| LOC102653738 | AmOr135 | 99.723 | Or135 | Group13 | 959662 | 964627 | 4966 | + | XM_026444706.1 |
| LOC100576992-a | AmOr136 | 100 | Or136-a | Group12 | 832688 | 837877 | 5190 | + | GSAman00055 |
| LOC724202-a | AmOr136 | 100 | Or136-b | Group12 | 788086 | 792399 | 4314 | + | GSAman00046 |
| LOC100576522 | NA | 0 | Or13a | Group5 | 13179353 | 13183135 | 3783 | + | XM_026440708.1 |
| Or14 | AmOr14 | 99.739 | Or14 | Group2 | 10017779 | 10019474 | 1696 | + | XM_026446350.1 |
| LOC100576153 | AmOr140N | 100 | Or140N | Group11 | 14447675 | 14449052 | 1378 | + | XM_026443618.1 |
| LOC726459 | AmOr141 | 99.534 | Or141-a | Group15 | 7781687 | 7783890 | 2204 | - | XM_026445563.1 |
| LOC408517 | NA | 0 | Or141-b | Group15 | 7767022 | 7772438 | 5417 | - | GSAman00086 |
| LOC102654452 | AmOr142 | 84.293 | Or142 | Group2 | 6431271 | 6434577 | 3307 | + | XM_026439157.1 |
| LOC100578842 | AmOr144 | 99.732 | Or144 | Group9 | 9755888 | 9758887 | 3000 | - | XM_026442801.1 |

| | | | | | | | | | |
|---|---|---|---|---|---|---|---|---|---|
| LOC102656567-a | AmOr145 | 100 | Or145 | Group9 | 9752135 | 9755364 | 3230 | - | GSAman00028 |
| LOC107965011 | AmOr146 | 100 | Or146 | Group9 | 9747243 | 9749449 | 2207 | - | XM_026442751.1 |
| LOC726834-b | AmOr148 | 100 | Or148 | GroupUN2 | 145511 | 147957 | 2447 | - | XM_016917294.2 |
| LOC100579011-a | AmOr149CN | 99.51 | Or149 | Group2 | 2498281 | 2501333 | 3053 | + | XM_026439166.1 |
| LOC100577715 | AmOr15 | 94.774 | Or15 | Group2 | 10019570 | 10022150 | 2581 | + | XM_016918033.2 |
| LOC100579011-b | AmOr150C | 100 | Or150 | Group2 | 2502381 | 2505531 | 3151 | + | XM_026439165.1 |
| LOC100578724 | AmOr151 | 100 | Or151 | Group2 | 2506791 | 2509529 | 2739 | + | NM_001327950.1 |
| LOC102655559 | AmOr152 | 100 | Or152 | Group2 | 2510408 | 2513388 | 2981 | + | XM_016910995.2 |
| LOC102655285 | AmOr154 | 99.204 | Or154 | Group2 | 1086473 | 1091023 | 4551 | - | XM_026439052.1 |
| LOC102654782 | AmOr155C | 99.717 | Or155C | Group13 | 1992220 | 1995962 | 3743 | - | XM_026444728.1 |
| LOC102654530 | AmOr156 | 100 | Or156 | Group2 | 2449158 | 2455786 | 6629 | - | XM_016911002.2 |
| LOC102654216 | AmOr158F | 86.486 | Or158F | GroupUN2 | 128605 | 130993 | 2389 | - | XM_006571914.3 |
| LOC102656271 | AmOr159CN | 83.721 | Or159CN | Group13 | 1037155 | 1044048 | 6894 | + | XM_026444635.1 |
| LOC100577671 | AmOr16 | 100 | Or16 | Group2 | 10022808 | 10024303 | 1496 | + | XM_003250719.4 |
| LOC724763 | AmOr160 | 100 | Or160 | Group1 | 16995814 | 16999400 | 3587 | + | XM_001120660.5 |
| LOC100577955 | AmOr161 | 98.21 | Or161 | Group5 | 9317331 | 9320058 | 2728 | - | XM_016912359.2 |
| LOC107965761 | AmOr163 | 92.727 | Or163 | Group2 | 1094196 | 1101231 | 7036 | - | XM_026439049.1 |
| LOC107965760 | AmOr164F | 100 | Or164F | Group2 | 1069799 | 1075614 | 5816 | + | XM_026439143.1 |
| LOC102655218 | AmOr165C | 100 | Or165C | Group2 | 1154485 | 1157492 | 3008 | + | XM_026439056.1 |
| LOC102679224 | AmOr166C | 100 | Or166C | Group2 | 1176521 | 1186579 | 10059 | + | GSAman00022 |
| LOC724673 | AmOr167C | 100 | Or167PAR | Group2 | 1142518 | 1149401 | 6884 | + | GSAman00019 |
| LOC102654878 | AmOr168 | 99.732 | Or168 | Group7 | 8695778 | 8697622 | 1845 | - | XM_026441699.1 |
| LOC102654841 | AmOr169 | 100 | Or169 | Group7 | 8692263 | 8695056 | 2794 | - | XM_006564278.3 |
| LOC100577634 | AmOr17 | 98.005 | Or17 | Group2 | 10027136 | 10028918 | 1783 | + | XM_026439171.1 |
| Or170 | AmOr170 | 100 | Or170 | Group7 | 8688183 | 8690955 | 2773 | - | NM_001242993.1 |
| LOC100577938 | AmOr18 | 100 | Or18 | Group2 | 10029386 | 10031210 | 1825 | + | XM_003250678.4 |
| Or19 | AmOr19 | 100 | Or19 | Group2 | 10031932 | 10033502 | 1571 | + | NM_001242966.1 |
| Or2 | AmOr2 | 100 | Or2 | Group1 | 5723757 | 5749095 | 25339 | + | XM_006563204.3 |
| LOC100577590 | AmOr20 | 100 | Or20 | Group2 | 10033903 | 10036219 | 2317 | + | XM_003250717.3 |
| LOC725861 | AmOr22 | 100 | Or22 | Group2 | 10038531 | 10040643 | 2113 | + | XM_001121659.5 |
| LOC102655825 | AmOr23 | 100 | Or23 | Group2 | 10041889 | 10043794 | 1906 | + | XM_006565884.3 |
| LOC102655857 | AmOr24 | 100 | Or24 | Group2 | 10043939 | 10046008 | 2070 | + | XM_006565885.3 |
| Or25 | AmOr25 | 97.094 | Or25 | Group2 | 10046322 | 10048685 | 2364 | + | XM_016917965.2 |
| Or26 | AmOr26 | 100 | Or26 | Group2 | 10049035 | 10051106 | 2072 | + | NM_001242968.1 |
| Or27 | AmOr27 | 100 | Or27 | Group2 | 10051858 | 10053414 | 1557 | + | NM_001242969.1 |
| Or30-a | AmOr29 | 100 | Or29 | Group2 | 10057004 | 10059406 | 2403 | + | XM_006565827.3 |
| LOC100577787 | AmOr3 | 99.752 | Or3 | Group2 | 9980044 | 9982278 | 2235 | + | XM_003250721.4 |
| Or30-b | AmOr30 | 100 | Or30 | Group2 | 10060394 | 10062281 | 1888 | + | NM_001242970.1 |
| LOC726097 | AmOr31 | 100 | Or31 | Group2 | 10062610 | 10064555 | 1946 | + | XM_001121864.5 |
| LOC100577334 | AmOr33 | 100 | Or33 | Group2 | 10067275 | 10069679 | 2405 | + | XM_026439118.1 |
| LOC113218602 | AmOr34 | 98.5 | Or34 | Group2 | 10069965 | 10072033 | 2069 | + | XM_026439122.1 |
| Or35 | AmOr35 | 99.516 | Or35 | Group2 | 10072854 | 10074595 | 1742 | + | NM_001242971.1 |
| LOC100577226 | AmOr36 | 100 | Or36 | Group2 | 10074586 | 10076944 | 2359 | + | XM_003250709.4 |
| Or37 | AmOr37 | 99.742 | Or37 | Group2 | 10077675 | 10079910 | 2236 | + | XM_006565823.3 |
| LOC107963966 | AmOr38 | 99.017 | Or38 | Group2 | 10080641 | 10082956 | 2316 | + | XM_006565890.3 |
| LOC100577101 | AmOr39 | 100 | Or39 | Group2 | 10083616 | 10085910 | 2295 | + | XM_003250706.4 |
| Or4 | AmOr4 | 100 | Or4 | Group2 | 9982958 | 9984808 | 1851 | + | NM_001242958.1 |
| LOC100577068 | AmOr40 | 100 | Or40 | Group2 | 10086850 | 10089330 | 2481 | + | XM_016918050.2 |
| Or41 | AmOr42 | 100 | Or41 | Group2 | 10090900 | 10092877 | 1978 | + | NM_001242973.1 |
| LOC100576984 | AmOr43 | 100 | Or43 | Group2 | 10094267 | 10096818 | 2552 | + | XM_003250703.4 |
| LOC100576944 | AmOr44 | 100 | Or44 | Group2 | 10098208 | 10100642 | 2435 | + | XM_003250702.4 |

| | | | | | | | | | |
|---|---|---|---|---|---|---|---|---|---|
| LOC100576914 | AmOr45 | 100 | Or45 | Group2 | 10102314 | 10104414 | 2101 | + | XM_006565895.3 |
| LOC100576881 | AmOr46 | 100 | Or46 | Group2 | 10104829 | 10106838 | 2010 | + | XM_026439172.1 |
| LOC113218533 | AmOr47 | 99.754 | Or47 | Group2 | 10108355 | 10110552 | 2198 | + | XM_026438981.1 |
| LOC100576816 | AmOr48 | 100 | Or48 | Group2 | 10112338 | 10114588 | 2251 | + | XM_006565899.3 |
| LOC100576783 | AmOr49 | 100 | Or49 | Group2 | 10114981 | 10117008 | 2028 | + | XM_006565901.3 |
| LOC107966034 | NA | 0 | Or4-like | Group16 | 57762 | 60321 | 2560 | + | XM_026445752.1 |
| Or5 | AmOr5 | 100 | Or5 | Group2 | 9986998 | 9988716 | 1719 | + | NM_001242959.1 |
| Or50 | AmOr50 | 99.487 | Or50 | Group2 | 10117134 | 10119750 | 2617 | + | XM_026446347.1 |
| Or51 | AmOr51 | 98.533 | Or51 | Group2 | 10121395 | 10123538 | 2144 | + | XM_026446358.1 |
| Or52 | AmOr52 | 100 | Or52 | Group2 | 10123985 | 10125552 | 1568 | + | NM_001242977.1 |
| Or53 | AmOr53 | 100 | Or53 | Group2 | 10126414 | 10127954 | 1541 | + | NM_001242978.1 |
| LOC102656907 | AmOr54 | 100 | Or54 | Group2 | 10128418 | 10130332 | 1915 | + | XM_006565903.3 |
| Or55 | AmOr55 | 99.51 | Or55 | Group2 | 10130630 | 10133207 | 2578 | + | XM_026438980.1 |
| Or56 | AmOr56 | 100 | Or56 | Group2 | 10134234 | 10136939 | 2706 | + | NM_001242980.1 |
| Or57 | AmOr57 | 99.509 | Or57 | Group2 | 10139205 | 10142463 | 3259 | + | NM_001242981.1 |
| Or58 | AmOr58 | 99.756 | Or58 | Group2 | 10142917 | 10144562 | 1646 | + | NM_001242982.1 |
| LOC102653637 | AmOr59 | 100 | Or59 | Group2 | 10144891 | 10147131 | 2241 | + | XM_006565906.3 |
| LOC102655367 | AmOr6 | 99.746 | Or6 | Group2 | 9990191 | 9992151 | 1961 | + | XM_016918052.2 |
| LOC102653703 | AmOr60 | 99.756 | Or60 | Group2 | 10147668 | 10149735 | 2068 | + | XM_006565907.3 |
| LOC102655435 | AmOr61 | 100 | Or61 | Group2 | 10149997 | 10152787 | 2791 | + | XM_016918048.2 |
| Or63-b | AmOr62 | 100 | Or62 | Group15 | 6970551 | 6973016 | 2466 | - | GSAman00030 |
| Or63-a | AmOr62 | 99.652 | Or63 | Group15 | 6974317 | 6976596 | 2280 | - | GSAman00031 |
| LOC410603 | AmOr64 | 99.491 | Or64 | Group15 | 6967145 | 6969218 | 2074 | - | XM_394081.6 |
| LOC107963999 | AmOr65 | 99.746 | Or65 | Group15 | 6964546 | 6966223 | 1678 | - | XM_016916515.2 |
| LOC100578045 | AmOr67 | 100 | Or67 | Group15 | 6957264 | 6963155 | 5892 | - | XM_026445628.1 |
| LOC100578886 | AmOr68 | 98.945 | Or68 | Group13 | 4482130 | 4483929 | 1800 | + | GSAman00007 |
| LOC100578751-a | AmOr69 | 98.551 | Or69 | Group13 | 4484993 | 4486634 | 1642 | + | GSAman00024 |
| LOC102655434 | AmOr7 | 100 | Or7 | Group2 | 9993982 | 9995638 | 1657 | + | XM_006565881.3 |
| LOC100578751-b | AmOr70 | 100 | Or70 | Group13 | 4486997 | 4489629 | 2633 | + | GSAman00025 |
| LOC100578685 | AmOr71 | 99.73 | Or71 | Group13 | 4490327 | 4491960 | 1634 | + | XM_016915872.2 |
| LOC100578617 | AmOr72 | 100 | Or72 | Group13 | 4492569 | 4498706 | 6138 | + | XM_016915873.2 |
| LOC100577755 | AmOr74C | 99.476 | Or74C | Group12 | 783507 | 787232 | 3726 | + | XM_003249181.4 |
| LOC102653814 | AmOr75 | 100 | Or75 | Group13 | 996241 | 1001387 | 5147 | + | XM_016917288.2 |
| LOC724202-b | AmOr76 | 98.237 | Or76 | Group12 | 793409 | 797163 | 3755 | + | GSAman00049 |
| LOC724462-a | AmOr77C | 100 | Or77C | Group12 | 796973 | 801157 | 4185 | + | GSAman00042 |
| LOC724462-b | AmOr78 | 100 | Or78 | Group12 | 802725 | 805514 | 2790 | + | GSAman00040 |
| LOC100576282 | AmOr79 | 100 | Or79 | Group12 | 805681 | 809658 | 3978 | + | XM_006558326.3 |
| LOC102655553 | AmOr8 | 100 | Or8 | Group2 | 9996230 | 9998090 | 1861 | + | XM_006565882.3 |
| LOC724590-a | AmOr80 | 100 | Or80 | Group12 | 810659 | 814475 | 3817 | + | GSAman00033 |
| LOC724590-b | AmOr81 | 81.087 | Or81 | Group12 | 815368 | 818589 | 3222 | + | GSAman00034 |
| LOC100576246 | AmOr82 | 100 | Or82 | Group12 | 818972 | 823344 | 4373 | + | XM_003249220.4 |
| LOC100576212 | AmOr83F | 99.754 | Or83F | Group12 | 823326 | 828106 | 4781 | + | XM_003249219.4 |
| LOC100576992-b | AmOr85 | 99.689 | Or85 | Group12 | 839665 | 841335 | 1671 | + | GSAman00056 |
| LOC100577024 | AmOr86 | 100 | Or86 | Group12 | 841903 | 845355 | 3453 | + | XM_003249233.4 |
| LOC102656838 | AmOr87 | 100 | Or87 | Group12 | 845583 | 850848 | 5266 | + | XM_016915586.2 |
| LOC102654183 | AmOr88 | 93.176 | Or88 | Group12 | 850992 | 856224 | 5233 | + | XM_026444443.1 |
| LOC100576940-a | AmOr89 | 97.03 | Or89 | Group12 | 856316 | 858723 | 2408 | + | GSAman00058 |
| Or9 | AmOr9 | 100 | Or9 | Group2 | 10000277 | 10002223 | 1947 | + | NM_001242960.1 |
| LOC100576940-b | AmOr90 | 96.659 | Or90 | Group12 | 858910 | 863200 | 4291 | + | GSAman00065 |
| LOC724911 | AmOr91 | 100 | Or91 | Group12 | 863552 | 867237 | 3686 | + | XM_016915581.2 |
| LOC725052 | AmOr94 | 100 | Or94 | Group12 | 872181 | 875072 | 2892 | + | GSAman00018 |

| | | | | | | | | | |
|---|---|---|---|---|---|---|---|---|---|
| LOC100576839 | AmOr96 | 100 | Or96 | Group12 | 875161 | 882860 | 7700 | + | XM_003249230.4 |
| Or98 | AmOr98 | 100 | Or98 | Group11 | 11138454 | 11140441 | 1988 | - | NM_001242985.1 |
