## Supplemental Table 2 for "Singular Odorant Receptor Expression Orchestrated by Promoter Activation Specificity in *Apis Mellifera* Olfactory Sensory Neurons": Supplemental table 2.pdf

Supplemental table 2. Primers used in the study for RNA-FISH.

| Genes | Primer sequence (5'-3') |
|---|---|
| Or154 | TTGTTGTTGGCCACGTCCTTTCAATAGTACCAAAAATCAAATTT |
| | GAATGTCTGCGTGAGCATATTTGGTGCACTTTTACTTTGG |
| | TATTTGGCTGCCACTGCTTTTATGCTAGGACCTGCAATA |
| | GTATCTCTCCAATGTTTTCTCCATTTTCACTACATTGCGTGTT |
| Or163 | TTGTAGCTCTCACGGCTGGGATGTTAATGGCATTTTCTGC |
| | AGCTGCATATGATATGCTTTGGTATAATTTAAATCCGCAAGAT |
| | ATTATGTTGCTGCGAATTGCGAAGCATTTATTCTTTGTTATACAG |
| | TGACGCTTTCTGCTGAAACTTTTGCTAGTATGCAAAAAATTTC |
| Or128 | AATTGTAGGACTGTGGCCATACGACAACTCAATCTACGTA |
| | GTTGGCAATTATTGTATGCATACTCTCATTCTCTGTAAAT |
| | CGGTTGTATTCTTGTAGTTGCTTATCATTTAATGATAGCA |
| | CGATGATGAGTTCCTCCTTCTCCTATTGTGCTGTTATATA |
| Or129F | ACTGTTTTCATACACAGAACTATTGTTCGATAACATACGT |
| | GACATTCATTGCCGCACTATCGAAAATATCCATATAGCCT |
| | AATCTTCGGTTCTGCTCTCATGGTTATGTATCATTTAATG |
| | GTTCATGATGTTCAAAAGTTCAGTGGGTTGTGAATTACGA |
| | GGTTTATTCACTGCATCTTATGCAGGCTTTACATCGATGA |
| Or25 | CTTGTAGCATGTTTTTGATGGTGGAAATCAAGAAGGTATATAACAG |
| | CCATTTCCTTCGTCGAGTTTGATCGCTGATATACGTTACAGC |
| | CGGATATGAGTAAAGCTATAGATGACAGGATTGCAAATATTGTAATTC |

| | |
|---|---|
| | ATATAGGCGATGGGGTTGCCGAACAGTGTCAAAAAGTTG |
| Or26 | CTACGAAGACGAGAGTTTGAAGACAAGAATGAAGGCAATCG |
| | GTTTCATCGCGTGCATCGCCGCACTTTGCATGCATAGCGG |
| | ATGATTTGCTTGGTTGAATTGGTGGGATGCACAATGAATATG |
| | ATCACTGCTGGAAAACTCATACAAATTTCTATCGCGACTTTT |
| Or27 | CTAAAACCAATCGGTGCTTGGCCTTTGTTCTCTACCACCAC |
| | ATGTGGCAGCCTCGTGCGCCATCTTCATGCAAGGTGGCAT |
| | GCACAGTTGGAATTTTCAGTCTCGCGGCTGTTCTTGCCGC |
| | GCACGATGATCATATGTATAGTCGGTTATTATATTCTTACGGAATG |
| Or40 | GCCACTTCCATCGTCCACCACGAAATTCGAAAAGATAATGAC |
| | GAGCTTCATGCAAGTTAGTACAATTTGTTTCGGCATCGTAT |
| | CTAATCTAGAGCACAGTATCGTTCTTGGTTTCCAATTTCTGAC |
| | TGCTTAGCTATTGTATTCTCGTGGAATGGTCTGGACGTGA |
| Or39 | TTTCTGTTGGATTACCACGCTATTCGT |
| | AGCATATGGAAGTTGATTGGAAGGCG |
| | ACCAATCCAGGATACGGTATTATCCTT |
| | ATTTGCATGTCACACTATTGGTCAGTT |
| Or169 | TGTCGAGGGAAAGAGCGTTGAAGGACAGCACGGTGAACAT |
| | TCGGTTCGCTTATGATTTGGGCATCGTTGCCTCTCGCTAA |
| | TACGCGACGTGACGATTGGTACGGTGAAACCGTTGGAGAA |
| | TTTCCAATCGGTGTTTTCATTGGCGGAGTTGCCCTACAGA |
| Or170 | AAGCGTTGAAGGATGGGACGGTGAACATAGCCATCTCGTT |

| | |
|---|---|
| | GCGAAACGACAATGTCAGACAGGAAATGAAGTTGGTAAC |
| | ATTTGAAGCGAGTTCATCTATAGCGGATGATGTGGTGTATG |
| Or62 | GCCTTCCATTCATTCTGGTACCAATCTCGAGGACGCA |
| | ATGTACGGTCCCTAGGACTCGTCTCCAATTTCACTTCTGC |
| | TGCTACTTACGACTACTGGAAATCTAGGTAGTGATTCGTTGT |
| | CTTTGCTCATACAGTTATATCTTTACGCCTACGCTGGCGAC |
| | GTCTTGGTAATTCGATCTACTTCTGTACGTGGTACGATATGCC |
| Or64 | CTTTTCATGACTGTTCCTATGCTTGCTGGAGATGAAGAAGAAG |
| | ACTCAAACTGAAGATCTTGGTAATTCGGTGTACTTCTGTACTT |
| | GCGACATGTTGAACGAAACAATTAGCTCTATTTTGATCGTACAG |
| Or65 | GGGAAAAGTGATGAGTTTTCGTGTACGTTCCACAGGACTTG |
| | CATGGGTAGAGTTGCATGCACTAGTTTAATATCTTGTTCGT |
| | CTTCTGCGGTTAACGATTACAACGAATCAAACGATGAAGAAAA |
| | TGGGGACGAAATACAAGTGATTAATGCAACGGAAGAAAATG |
| | AGCATTGGAAATATTGTCTTGACGATAAAAATACTTATAATAATGTGC |
| Or67 | ATACGTCCTGTAGGACTCGTTTCCAATTTTTCTTCTGCGAT |
| | CTCGTTTATCGCGTCCACGCTTTTCATGACTGTGCCTA |
| | AAAATACTTGCGGAGATATCCATTCTTTTGATACAATTGTTCGC |
