## Supplemental Table 3 for "Singular Odorant Receptor Expression Orchestrated by Promoter Activation Specificity in *Apis Mellifera* Olfactory Sensory Neurons": Supplemental table 3.pdf

Supplemental table 3. Primers used in the study for PCR.

| Genes | Primer sequence (5'-3') |
|---|---|
| Or25-Or26 | F1: CTTTGCTGCATTTTGATCATTGCAATG |
| | R1: GAATGGACAAGGTAACCGATACATCG |
| Or26-Or27 | F2: GGCAAAAAAGTGGGAGAAAAATTTTATATGACTG |
| | R2: TAATTTCTTCTGTGATCGCGGTTTGGTTCATC |
| Or39-Or40 | F: TGACTGCCGGAAAGATAATGGATATGTC |
| | R: CCGAACTTTTTCTGAATTGCGGTAATATCA |
| Or169-Or170 | F: GTACACGAAAAATGGTATCTTCATGACGTAAAGTTTCAACATATA |
| | R: CAATTCTATTATCCAAGCGACGACGAAATAGATTTTC |
